# The BMP ligand Gdf6a Regulates Development of the Zebrafish Craniofacial Skeleton in a Pharyngeal Arch-Specific Manner

**DOI:** 10.1101/2025.11.14.688493

**Authors:** Sabrina C. Fox, Sarah M. Hay, Andrew J. Waskiewicz, Jennifer C. Hocking

**Affiliations:** Department of Biological Sciences, University of Alberta, Edmonton, Alberta, Canada; Women & Children’s Health Research Institute, University of Alberta, Edmonton, Alberta, Canada; Division of Anatomy, Department of Surgery, University of Alberta, Edmonton, Alberta, Canada; Department of Cell Biology, University of Alberta, Edmonton, Alberta, Canada; Department of Medical Genetics, University of Alberta, Edmonton, Alberta, Canada

**Keywords:** BMP signaling, craniofacial development, zebrafish, cartilage, chondrogenesis

## Abstract

Craniofacial development involves the concerted action of several different tissue types and cellular processes that must be intricately regulated. Here, we describe the role of the BMP ligand Growth and Differentiation Factor 6a (*gdf6a*) in the development of the zebrafish craniofacial skeleton. Larval *gdf6a* mutant zebrafish have malformations in the midline of the craniofacial skeleton that correlate with the expression of *gdf6a* in the embryonic pharyngeal arches. We show that Gdf6a has arch-specific roles in craniofacial morphogenesis; Gdf6a promotes chondrogenesis and alignment of midline craniofacial elements in the posterior pharyngeal arches (arches 2 and 3) and regulates morphogenesis within the mandibular symphysis of pharyngeal arch 1. We demonstrate that Gdf6a regulates craniofacial development through activation of canonical BMP signaling, likely acting cooperatively with additional BMP ligands. Taken together, this work elucidates how Gdf6a*/*BMP signaling directs development of the craniofacial skeleton, specifically along the ventral midline.

**Summary Statement:** Gdf6a regulates the development of the midline structures in the zebrafish craniofacial skeleton in an arch-specific manner via canonical BMP signaling.

## Introduction

Development of the craniofacial skeleton, which encompasses the neurocranium and viscerocranium, is a highly complex process (G. H. . Sperber et al., 2010). In all vertebrates, the craniofacial skeleton is formed during embryonic development from cranial neural crest (Jiang et al., 2002; Lièvre & Douarin, 1975; Noden, 1978; Yoshida et al., 2008). After delamination from the neural tube, neural crest cells migrate ventrally through the head mesenchyme and populate the frontonasal process and the pharyngeal arches, where they differentiate into the connective tissues of the face and neck (Lièvre & Douarin, 1975; Noden, 1978). Aberrant development can result in craniofacial abnormalities, which significantly impair the functioning and quality of life of affected individuals (Gorlin et al., 2001). Indeed, understanding the factors that regulate craniofacial development could provide insight into the pathogenesis and etiology of these abnormalities in human patients, allowing for proper counselling and possible therapeutic intervention.

Craniofacial development involves the coordination of many cellular and developmental processes that require extensive spatial and temporal regulation. One critical regulator is the Bone Morphogenetic Protein (BMP) signaling pathway. BMP signaling is initiated when a ligand dimer binds to its cognate type I and type II receptors (Yamashita, ten Dijke, et al., 1994). Ligand/receptor interactions trigger the phosphorylation of receptor-associated Smad (r-Smad) proteins, which then form a complex with the common Smad (co-Smad) Smad4 and enter the nucleus, where they bind to DNA motifs termed BMP response elements (BREs) and regulate the expression of target genes (Katagiri et al., 2002; Kawabata et al., 1998). Moreover, several negative regulators of the BMP signaling pathway exist, allowing for precise temporal and spatial regulation of BMP signaling. These negative regulators include secreted extracellular antagonists such as Noggin (NOG) and BMP and Activin Membrane-Bound Inhibitor (BAMBI), which block ligand-receptor interactions, and inhibitory Smads (i-Smads) such as Smad6, which inhibit the pathway intracellularly by interfering with r-Smad activity (Groppe et al., 2002; Imamura et al., 1997; Nakao et al., 1997; Onichtchouk et al., 1999; Zimmerman et al., 1996).Notably, at least 20 ligands belong to the BMP family, and overlapping and compensatory activity may occur among several members, further adding complexity to the dynamics of this signaling pathway (Ducy & Karsenty, 2000).

One member of the BMP signaling pathway important for skeletal development is Growth and Differentiation Factor 6 (*GDF6*). *GDF6* shares a high degree of sequence similarity with *GDF5* and *GDF7*, thereby forming a unique subgroup of BMP ligands (Chang et al., 1994; Storm et al., 1994)*. Gdf5* and *6* are expressed in the presumptive joints in the murine appendicular skeleton (Settle et al., 2003; Wolfman et al., 1997),and *Gdf5^-/-^;Gdf6^-/-^* mutant mice display loss of joints that primarily reside in the distal appendicular skeleton (Settle et al., 2003). Furthermore, mutations in *Noggin,* which encodes a secreted antagonist that binds to Gdf5 and Gdf6, causes loss of all joints, indicating that regulation of BMP signaling is critical for proper joint development (Brunet et al., 1998; Miyamoto et al., 2007; Valdes et al., 2011).

*GDF6* is also implicated as a critical regulator of craniofacial development. *Gdf6^-/-^* mutant mice display alterations to the craniofacial skeleton, including premature cranial suture closure and fused inner ear ossicles (Clendenning & Mortlock, 2012; Settle et al., 2003). Furthermore, several instances of craniofacial abnormalities in patients with mutations in *GDF6,* variants in *GDF6* enhancers, or chromosomal abnormalities encompassing *GDF6* have been reported, including cleft palate, laryngeal cartilage abnormalities leading to vocal impairment, facial asymmetry, and hearing loss due ossicle fusion (Clarke et al., 1994, 1995, 1998; McGaughran et al., 1998). Despite evidence from mouse and human research implicating Gdf6/GDF6 as a key regulator of craniofacial development, little is known about the particular functions of Gdf6/GDF6 in this process.

In recent years, zebrafish has become a common tool for studying the molecular mechanisms underlying craniofacial development (Mork & Crump, 2015). Many mammalian craniofacial structures, including the upper and lower jaws, palate, hyoid bone, ossicles, and laryngeal cartilages, have direct homologs in zebrafish, thereby allowing for the study of their formation in the zebrafish (Mork & Crump, 2015). The cartilage template for the zebrafish craniofacial skeleton is fully developed by 5 days post fertilization (dpf), with ossification starting as early as 4 dpf, allowing for rapid analysis of skeletal and craniofacial phenotypes (Cubbage & Mabee, 1996). Furthermore, several resources for studying both BMP signaling and craniofacial development have been developed in zebrafish, including transgenic strains that provide readouts of signaling activity or label cell types and mutant strains to probe gene function (Alexander et al., 2011; Dutton et al., 2008a). Indeed, zebrafish are a model well-suited for studying the intersection of BMP signaling and craniofacial development.

To investigate the etiology of craniofacial abnormalities in patients with disrupted GDF6 signaling, we analyzed the role of the zebrafish homolog of *GDF6/Gdf6, gdf6a,* in craniofacial development. We observe *gdf6a* mRNA expression in the pharyngeal arches from 48-72 hours post fertilization, stages that correspond to morphogenesis of the craniofacial skeleton, in regions that correspond to cartilage elements that lie along the ventral midline of the craniofacial skeleton. Accordingly, we report that several midline craniofacial cartilages are malformed or absent in *gdf6a^-/-^* mutants. We show that BMP signaling is active in these regions and is necessary for the development of midline craniofacial cartilages, and that BMP signaling via Gdf6a is necessary for proper craniofacial development. Mechanistically, we observe arch-specific roles for Gdf6a in craniofacial skeletal development; in the posterior arches, *gdf6a* promotes the formation of the ceratohyal and hypobranchial cartilages by activating a chondrogenic program, whereas in the mandibular arch, Gdf6a regulates the morphogenesis of the mandibular symphysis, a structure that connects the bilateral arms of Meckel’s cartilage. Taken together, this work provides insight into the role of Gdf6a/Gdf6/GDF6 in development of the vertebrate craniofacial skeleton.

## Results

### *gdf6a* is expressed in the pharyngeal arches during craniofacial development

To gain insight into the potential role of Gdf6a in skeletal development, we analyzed the expression of *gdf6a* mRNA using whole mount *in-situ* hybridization at several stages of zebrafish embryonic development. At the pharyngula stage (24 and 36 hours post fertilization (hpf)), *gdf6a* is localized primarily to the dorsal eye and epidermis, where it has previously documented roles (Figure S1) (French et al., 2009; Gosse & Baier, 2009; Gramann et al., 2019). *gdf6a* is also expressed weakly in the ventral head and otic vesicle at 36 hpf (Figure S1). By 48 hpf, expression has increased dramatically in the otic vesicle and can be clearly observed in the perioral region anterior to the stomodeum (presumptive mouth) and ventral domain of pharyngeal arches 1 and 2, with weaker expression in posterior arches (Figure 1A-A’, Figure S1). At 60 and 72 hpf, *gdf6a* mRNA is expressed in a segmental pattern in the ventral domain of each pharyngeal arch; more specifically, along the midline of the presumptive Meckel’s, ceratohyal, and hypobranchial cartilages (Figure 1B-C’, Figure S1). As the cartilages appear by 72 hpf, *gdf6a* expression in the medial pharyngeal arches from 48-72 hpf is well placed to regulate development of the ventromedial craniofacial skeleton.

**Figure 1.**
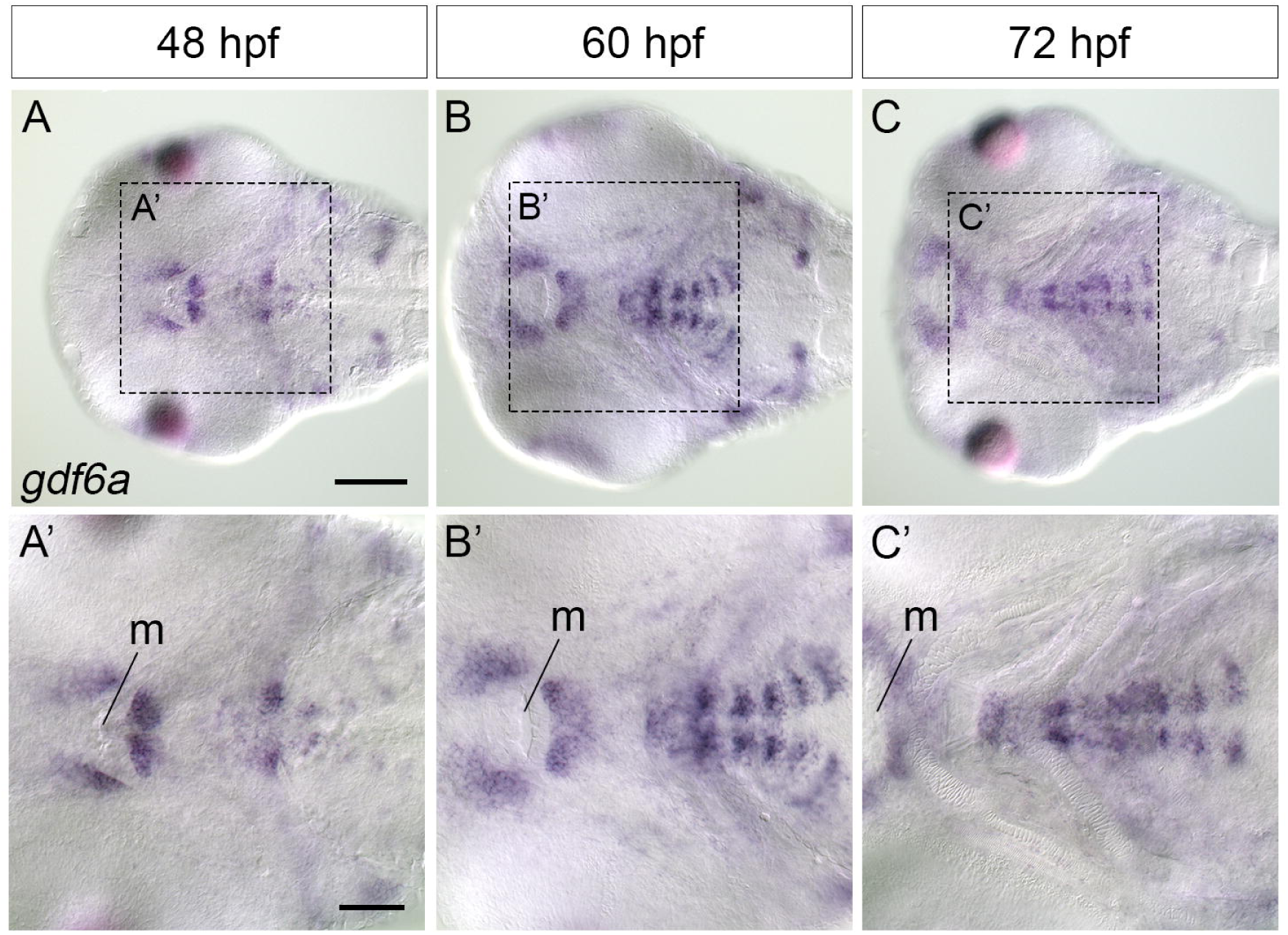
*gdf6a* is expressed in the pharyngeal arches. (A-C’) DIC microscopy images of zebrafish showing *gdf6a* mRNA expression from 48-72 hpf. Expression increases over time, but at all three stages *gdf6a* is largely localized to the midline in the ventral head. m=mouth, pas=pharyngeal arches. Scale bars=100 microns in A, 50 microns in A’.

*GDF6,* the human paralog of *gdf6a,* is frequently grouped with *GDF5* and *GDF7* to form the *GDF5/6/7* subfamily based on sequence similarity and evolutionary relatedness (Wolfman et al., 1997). Indeed, like the human paralogs *GDF5/6/7,* the zebrafish paralogs *gdf5/6a/6b/7* share considerable sequence similarity in the C-terminal region that gives rise to the mature signaling peptide, suggesting potential overlapping activity (Figure S2). To determine whether zebrafish *gdf5, gdf6b,* and *gdf7* are active in the pharyngeal arches, we performed *in-situ* hybridization at 60 hpf (Figure S3). Although all are somewhat expressed in the pharyngeal arches, only *gdf5* shares any spatial overlap with *gdf6a* in the pharyngeal skeleton, and only in the second arch at the presumptive basihyal and ceratohyal joint (Figure S3). Therefore, *gdf6a* has a unique expression domain during craniofacial development relative to its family members.

### *gdf6a^-/-^* mutants have craniofacial malformations

To examine the function of Gdf6a in formation of the craniofacial skeleton, we performed phenotypic characterization of *gdf6a^-/-^* mutants at 5 days post fertilization (dpf), by which time the cartilage elements of the larval viscerocranium have formed (Cubbage & Mabee, 1996). Stereomicroscope images of live *gdf6a^-/-^* mutants and siblings (notated as *gdf6a^+/?^*) confirmed prior findings that *gdf6a^-/-^* mutants have severely reduced eye size and enhanced pigment formation compared to siblings, but also highlighted that *gdf6a^-/-^* mutants have altered head shape; while siblings have a “tapered” head shape, *gdf6a*^-/-^ heads are blunted and rounder, suggesting the craniofacial skeleton is morphologically altered (French et al., 2009; Gosse & Baier, 2009; Gramann et al., 2019) (Figure 2A, B). Views of larvae stained with Alcian blue and Alizarin red to label cartilage and bone, respectively, revealed a reduction in overall size of the craniofacial skeleton is *gdf6a^-/-^* mutants, and alterations to the midline of several cartilage elements (Figure 2C-F). In DIC images of flatmounted larvae, we observed abnormal morphology of Meckel’s and ceratohyal cartilages in *gdf6a^-/-^* mutants. Specifically, the cells in the mandibular symphysis were disorganized, resulting in a thickening of the joint (Figure 2G-H, M). Similarly, the ceratohyals, which articulate along the ventral midline of the zebrafish craniofacial skeleton, are misoriented in *gdf6a^-/-^* mutants (Figure 2G-H, arrowhead), resulting in a “splayed” appearance (Figure 2N). The basihyals, which run along the midline of the viscerocranium, and the hypobranchials, which connect the ceratobranchials to the basihyal, are both severely reduced in mutants (Figure 2I-J, O, P). In summary, *gdf6a^-/-^* mutants display malformation of midline structures within the developing viscerocranium.

**Figure 2.**
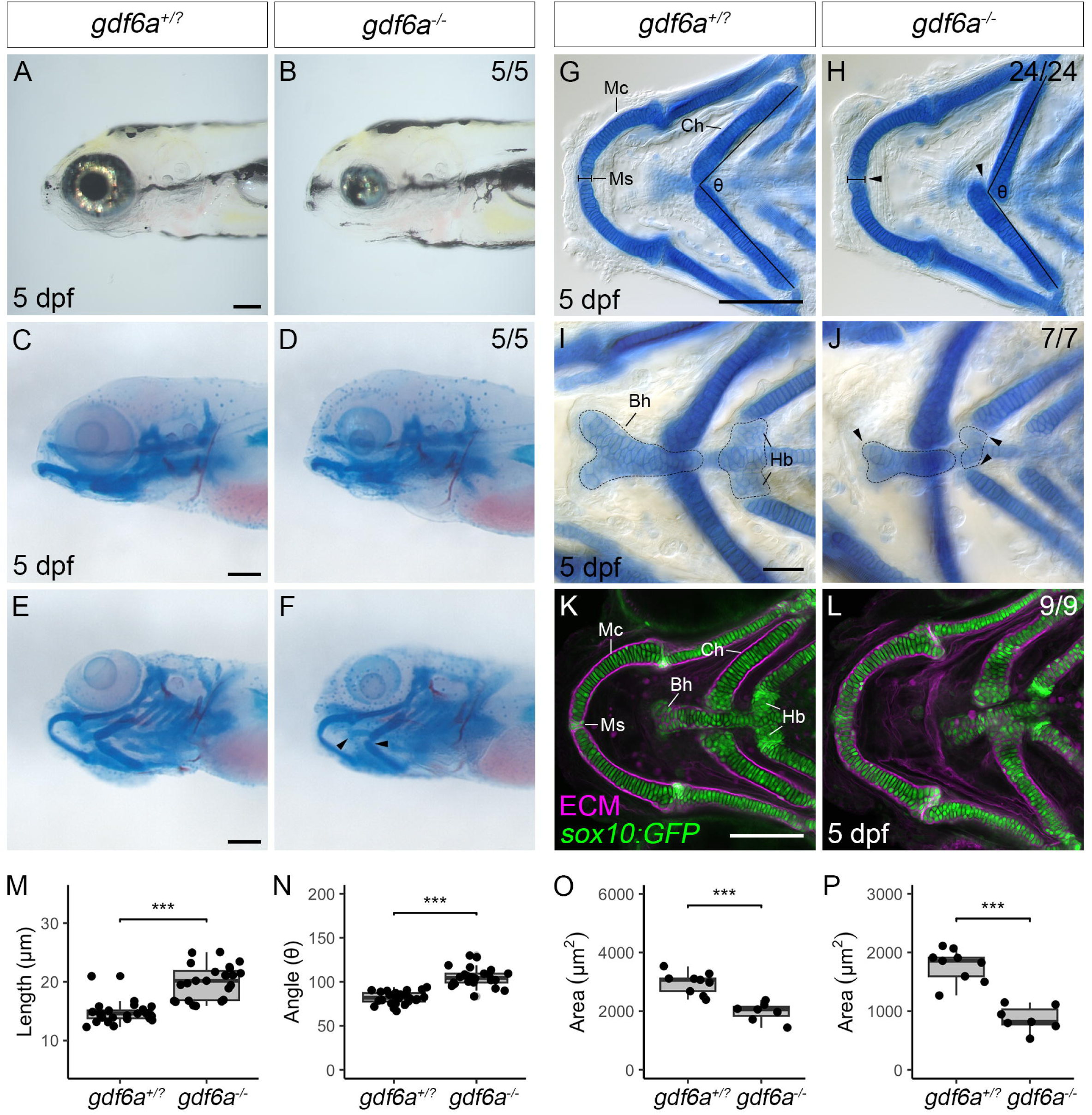
*gdf6a^-/-^* mutants have a craniofacial phenotype. (A-B) Stereomicroscope images of live *gdf6a^+/?^* and *gdf6a^-/-^* larvae at 5 dpf. (C-F) Stereomicroscope images of *gdf6a^+/?^* and *gdf6a^-/-^* larvae stained with Alcian blue (cartilage) and Alizarin red (bone). *gdf6a^-/-^* mutants have less tapered heads than siblings but overall structure of the craniofacial skeleton is maintained. (G-J) DIC microscopy images of ventral flatmounts of the craniofacial skeletons from *gdf6a^+/?^* and *gdf6a^-/-^* larvae stained with Alcian blue and Alizarin red at 5 dpf. Ceratohyals are misoriented along the midline and the cells of the mandibular symphysis appear disorganized in *gdf6a^-/-^* mutants compared to siblings (H, arrowheads). The quantified width of the mandibular symphysis and the angle of the ceratohyal are annotated in each panel. (I, J) Increased magnification reveals that the basihyal is reduced in size and the hypobranchials are almost completely absent in *gdf6a^-/-^* mutants compared to siblings (J, arrowheads). Dotted lines indicate the area of the basihyal and hypobranchials that were quantified. (K-L) Confocal microscopy images of *gdf6a^+/?^*(K) and *gdf6a^-/-^* (L) larvae on a *Tg(sox10:GFP)* transgenic background (green) and counterstained with wheat germ agglutinin (magenta) to visualize cell morphology. The disorganization of the mandibular symphysis, reduction in the basihyal, and absence of the hypobranchials in *gdf6a^-/-^* mutants is apparent. (M-P) Quantification of mandibular symphysis width (M), angle created by the ceratohyals (N), size of the basihyal (O) and hypobranchial 3 (P). All data analyzed by unpaired t-test. *** = p<0.001. Mc=Meckel’s cartilage, Ms=mandibular symphysis, Ch=ceratohyal, Bh=basihyal, Hb=hypobranchials, ECM=extracellular matrix. Scale bars=100 microns in A, C, G and K and 50 microns in I.

In addition to defects in the viscerocranium, we noted changes to nearby regions of the neurocranium of *gdf6a-/-* mutants. While the overall shape of the ethmoid plate (the zebrafish equivalent of the mammalian palate) is comparable between siblings and *gdf6a^-/-^* mutants, plate size is mildly reduced and the rods of cartilage that connect the ethmoid plate to the skull vault known as the trabeculae are malformed in *gdf6a^-/-^* mutants; rather than being continuous rods of cartilage, the trabeculae in *gdf6a^-/-^* mutants frequently have gaps such that the trabeculae no longer attach to the skull (Figure S4). 21% of larvae have unilateral gaps in the trabeculae, 5% have bilateral gaps, and 63%have no gaps but trabeculae that are bent compared to *gdf6a^+/?^* siblings (Figure S3). Notably, *gdf6a* is expressed in the maxillary domain at 36 hpf (Figure S3), a region containing neural crest cells that give rise to the ethmoid plate and trabeculae.

Next, facial musculature was visualized in 96 hpf *gdf6a^-/-^* mutants and siblings through staining with the F-actin label phalloidin. In ∼45% of *gdf6a^-/-^* mutants, at least one muscle fiber is mislocalized compared to siblings (Figure S5). This mislocalization is confined to muscle fibers attached to the midline of the larval craniofacial skeleton, including the intramandibularis anterior, intramandibularis posterior, interhyoides, and hyohyiodes (Figure S5). In contrast, muscles that lie more laterally in the craniofacial skeleton are unaffected. Given that muscles are attached to cartilage via tendons, we visualized craniofacial tendons using Thrombospondin 4 (Tbs4) immunofluorescence (Subramanian & Schilling, 2014). Unexpectedly, craniofacial tendons form normally in *gdf6a^-/-^* mutants except for subtle disruption of the sternohyoideus tendon (Figure S5). Further, the spatial expression of mRNA encoding several tendon markers, including *xirp2a, scxa*, *tnmd,* and *col1a2*, is unchanged in *gdf6a^-/-^* mutants (Figure S5). Taken together, this data indicates that Gdf6a regulates the attachment of muscles to cartilage independently of tendon formation in the zebrafish larval craniofacial skeleton.

### *gdf6a* regulates chondrogenesis of the hypobranchials

During craniofacial development, neural crest cells in each pharyngeal arch will give rise to structures that are anatomically and functionally distinct (Schilling & Kimmel, 1994). Therefore, the cell behaviors and molecular cues present in each arch are unique, and we hypothesized that Gdf6a has distinct roles in the formation of the mandibular, hyoid, and branchial arch structures. To test this, we fixed *gdf6a^-/-^* mutants and siblings carrying the *sox10:GFP* transgene at various stages of craniofacial development and imaged the developing craniofacial skeleton at several axial levels. We first focused on the z-stacks that illustrated development of derivatives of pharyngeal arch 3 (the first branchial arch). Mutant embryos fixed at a relatively early stage of development (54 hpf) have craniofacial anatomy indistinguishable from sibling controls (Figure 3A, B). At 60 hpf, we observe a lack of *sox10:GFP*+ cells in the region corresponding to the arch 3 hypobranchials (Figure 3C, D). As development proceeds, presumptive chondrocytes continue to be recruited to the hypobranchials in *gdf6a^+/?^* siblings and yet fail to form in mutants (Figure 6E, F, Figure S6). By 84 hpf, the hypobranchials are dramatically reduced in *gdf6a^-/-^* mutants compared to the well-formed hypobranchials in siblings (Figure S6).

**Figure 3.**
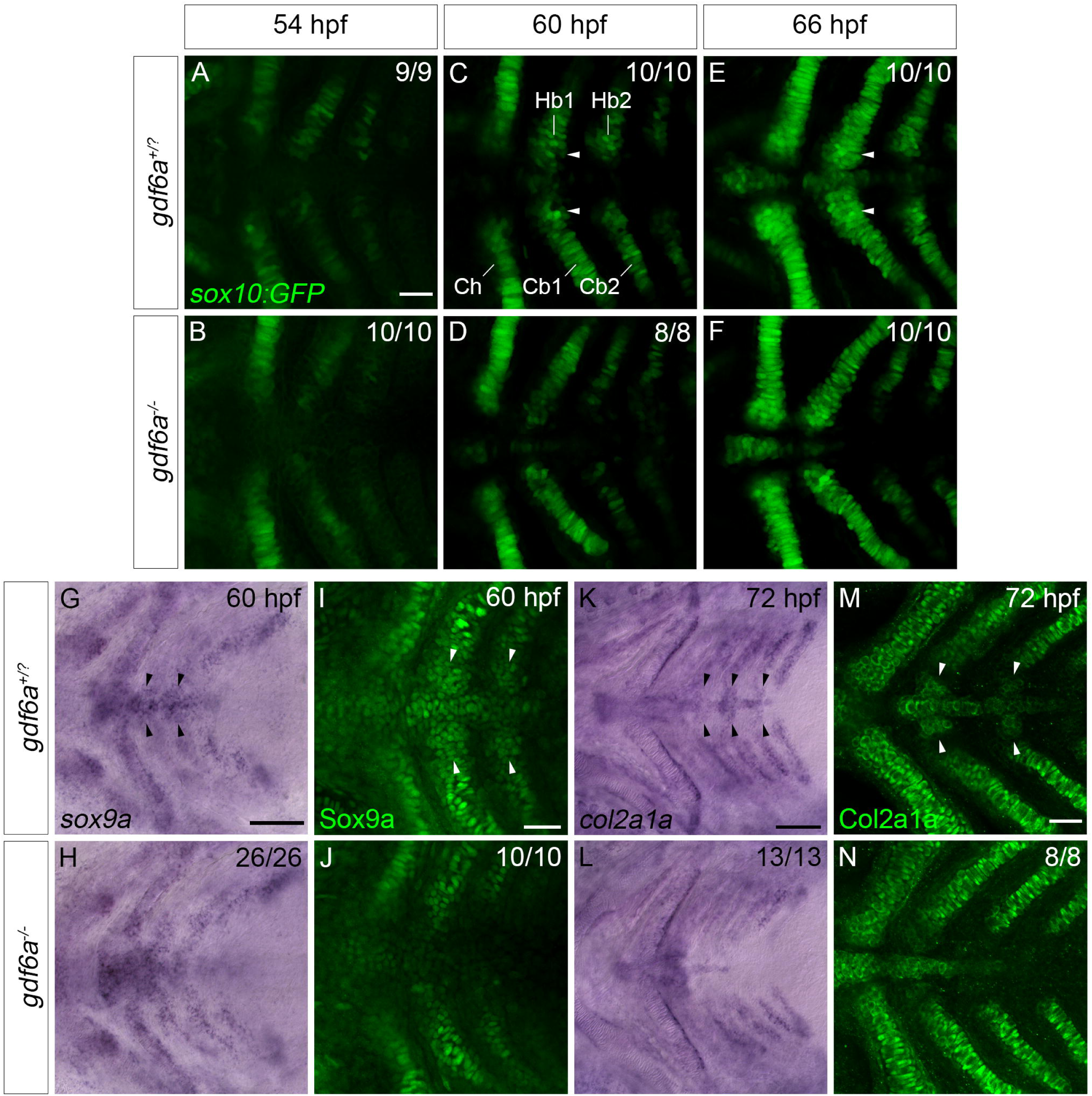
*gdf6a* regulates formation of the hypobranchials. (A-F) Single confocal z-slices of the posterior pharyngeal arches of *gdf6a^+/?^;Tg(sox10:GFP)* and *gdf6a^-/-^;Tg(sox10:GFP)* zebrafish at 54, 60, and 66 hpf. At 54 hpf, the pharyngeal cartilage has just condensed and started to differentiate. Hypobranchials (arrowheads, C and E) are largely absent in *gdf6a^-/-^* mutants from 60 hpf. (G-H) DIC microscopy photos of zebrafish embryos at 60 hpf showing *sox9a* mRNA expression in the posterior pharyngeal arches. (I-J) Single confocal z-slices of the posterior pharyngeal arches at 60 hpf showing Sox9a protein localization visualized with immunofluorescence. (K-L) DIC microscopy photos of zebrafish embryos at 72 hpf showing *col2a1a* mRNA expression in the posterior pharyngeal arches. (M-N) Single confocal z-slices of the posterior pharyngeal arches at 72 hpf showing the localization of Type II collagen (Col2a1a) immunofluorescence. Arrowheads in G, I, K, M point to hypobranchials, which are absent in *gdf6a^-/-^* mutants. Ch=ceratohyal, Hb1=hypobranchial 1, Hb2=hypobranchial 2, Cb1=ceratobranchial 1, Cb2=ceratobranchial 2. Scale bars=50 microns.

Consistent with a loss of hypobranchial chondrogenic progenitors, mRNA and protein expression for the chondrogenic transcription factor Sox9a and collagen protein Col2a1a are reduced along the midline in the posterior arches of *gdf6a^-/-^* mutants at 60 hpf (Figure 7G-N, arrowheads).

To determine whether the absence of hypobranchial chondrogenic progenitors is a consequence of decreased cell division or increased apoptosis, we performed immunofluorescence for phosphorylated Histone H3 (pH3) to mark M-phase cells and cleaved Caspase 3 to mark apoptotic cells at 54, 60, and 72 hpf. While there is considerable pH3 and Caspase 3 signal in the lateral regions of *gdf6a^+/?^* siblings at all stages observed, there is very little signal in regions closer to the midline of the posterior pharyngeal arches, suggesting that neither cell division nor cell death are prominent drivers of hypobranchial formation (Figure S7). Accordingly, we observed no obvious increase or decrease in either pH3+ or Caspase 3+ cells in the midline of the posterior pharyngeal arches in *gdf6a^-/-^* mutants at all stages examined (Figure S7), indicating that Gdf6a regulates chondrogenesis of the hypobranchials independently of cell division or death.

### *gdf6a* regulates chondrogenesis in pharyngeal arch 2

To assess Gdf6a’s role in mandibular and hyoid arch development, we analyzed confocal z-stacks of the craniofacial skeleton in the *Tg(sox10:GFP)* larvae at various timepoints. There is little morphological difference in the mandibular symphysis and ceratohyal joint between *gdf6a^+/?^* siblings and *gdf6a^-/-^* mutants at 54, 60 and 66 hpf (Figure S8). However, by 72 hpf, differences between mutant and sibling embryos become apparent. At 72 hpf, control fish have a “bridge” of chondrocytes along the midline that connects the ceratohyals (Figure 4A, Figure S8). In *gdf6a^-/-^* animals, this chondrocyte bridge is missing (Figure 4B, Figure S8). As development proceeds, the ceratohyals become progressively more misoriented in *gdf6a^-/-^* mutants (Figure 4C-J, Figure S8), indicating that the chondrocyte bridge connecting each arm of the ceratohyal to the midline is necessary for proper orientation of the ceratohyals.

**Figure 4.**
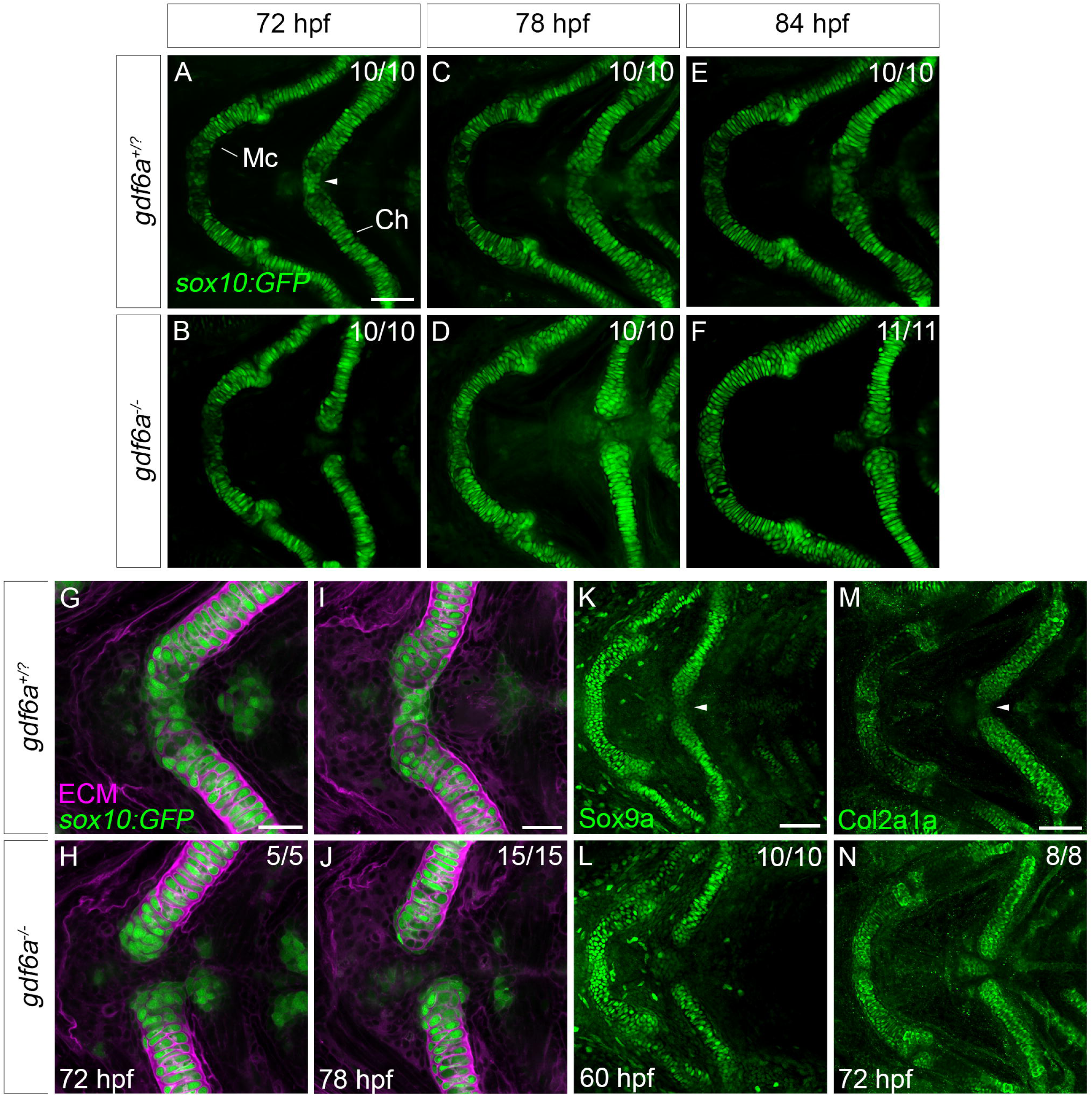
*gdf6a* regulates chondrogenesis of the ceratohyal joint. (A-F) Single confocal z-slices of the mandibular and hyoid arches of *gdf6a^+/?^;Tg(sox10:GFP)* and *gdf6a^-/-^; Tg(sox10:GFP)* zebrafish at 72, 78, and 84 hpf. In *gdf6a^+/?^* siblings, there is a bridge of chondrocytes at the ceratohyal joint that connects the bilateral arms of the ceratohyal (A, arrowhead); this chondrocyte bridge is missing in *gdf6a^-/-^* animals and ceratohyals become progressively more misaligned. (G-J) Single confocal z-slices of the ceratohyal joint from *gdf6a^+/?^;sox10:GFP* and *gdf6a^-/-^;sox10:GFP* animals with ECM visualized with wheat germ agglutinin. (K-N) Single confocal z-slices of *gdf6a^+/?^* siblings and *gdf6a^-/-^* mutants immunolabelled for Sox9a or Col2a1a. Arrowheads in K and M point to labelling of chondrocyte bridge present in *gdf6a^+/?^* siblings and absent in *gdf6a^-/-^* mutants. Mc=Meckel’s cartilage, Ch=ceratohyal. Scale bars=50 microns in A, K, M, 25 microns in G, I.

To test whether the absence of the cells that connect the ceratohyals results from a lack of chondrogenic progenitors in this region, we conducted Sox9s and Col2a1a immunofluorescence on *gdf6a^-/-^* mutants and *gdf6a^+/?^* siblings at 60 and 72 hpf, respectively. At 60 hpf, a small population of Sox9a+ cells flanked by the ceratohyals is visible in *gdf6a^+/?^* siblings (Figure 4K, arrowhead) but absent in *gdf6a^-/-^* mutants (Figure 4L). Further, we observed a reduction of Col2a1a protein localization at the region between the ceratohyals in *gdf6a^-/-^* mutants (Figure 4M) compared to *gdf6a^+/?^* siblings (Figure 4N, arrowhead), indicating that chondrogenesis is impaired in this region. Interestingly, we observed no change in Sox9a or Col2a1a protein expression in the mandibular symphysis in *gdf6a^-/-^* mutants of either age (Figure 4K-N). Therefore, we conclude that Gdf6a regulates the induction of chondrogenesis in the ventral hyoid arch from 60-72 hpf but has little effect on chondrogenesis of the mandibular symphysis.

### *gdf6a* regulates morphogenesis of the mandibular symphysis

We had initially observed a thickening of the mandibular symphysis in 5 dpf *gdf6a^-/-^* mutants, suggesting an error in development despite normal chondrogenesis. Notably, the morphology and extracellular matrix (ECM) composition of the mandibular symphysis is distinct to that of the rest of Meckel’s cartilage; cells that reside in the symphysis are rounder and secrete less ECM, as marked by WGA staining and Col2a1a immunofluorescence, compared to cells that reside more laterally in Meckel’s cartilage (Figure 5A-B’). Moreover, readouts of BMP signaling, including pSmad1/5 protein localization and *BRE:GFP* transgene expression, are localized to the round, low ECM-secreting cells, indicating that midline BMP signaling is appropriately localized to regulate the morphology and ECM secretion of these cells (Figure 5C-D’). Therefore, we hypothesized that Gdf6a regulates the morphogenesis of the mandibular symphysis. By imaging the mandibular symphyses of 72 hpf *gdf6a^-/-^* mutants and siblings labelled with the *sox10*:GFP transgene and WGA, we noted a disorganization of cells in the mutants (Figure 5E-F). At 78 hpf, cellular disorganization is especially apparent in the center of the mandibular symphysis where the bilateral arms of Meckel’s cartilage meet and results in a thickening of the mandibular symphysis in *gdf6a^-/-^* mutants (Figure 5G-I). Notably, the disorganization is not a result of disrupted planar cell polarity (PCP), as indicated by normal localization of γ-tubulin in the mutants (Figure S9). Taken together, this data indicates that, in contrast to the posterior arches, where Gdf6a induces chondrogenesis, Gdf6a regulates the morphogenesis of the mandibular symphysis joining the first arch cartilages.

**Figure 5.**
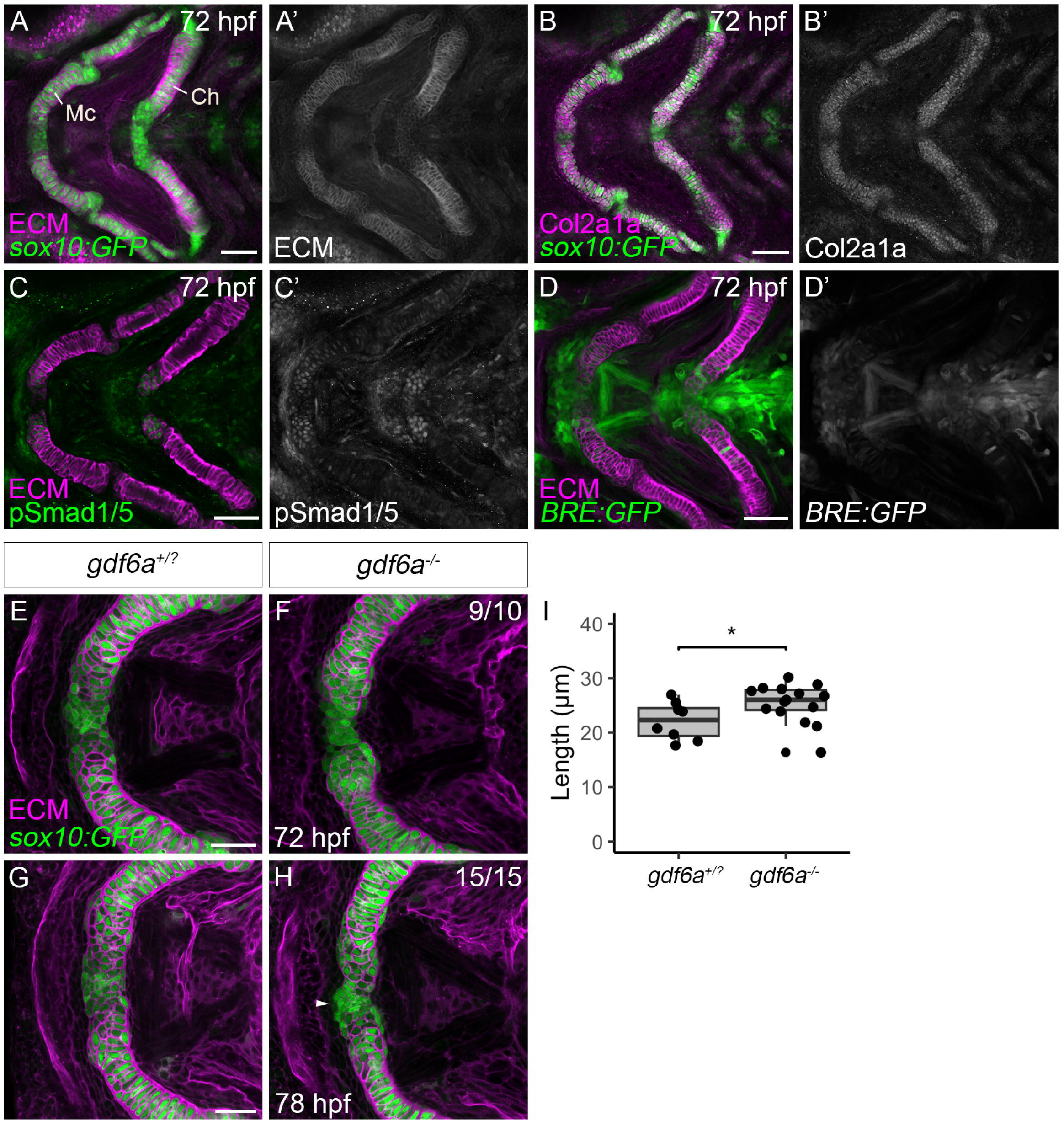
mandibular symphysis morphogenesis is disrupted in *gdf6a*^-/-^ mutants. (A-B’) Single confocal z-slices of wild type *Tg(sox10:GFP)* zebrafish at 72 hpf with extracellular matrix (ECM) or type II collagen (Col2a1a) visualized with wheat germ agglutinin staining or immunofluorescence, respectively. Cells in the mandibular symphysis and ceratohyal joint have distinct morphology and reduced ECM staining compared to cells in the arms of the ceratohyal and Meckel’s cartilage. (C-C’) Single confocal z-slices of wild type embryos at 72 hpf with labelled with phosphorylated Smad1/5 (pSmad1/5) antibody and wheat germ agglutinin. (D-D’) Single confocal z-slices of wild type *Tg(BRE:GFP)* embryos at 72 hpf with wheat germ agglutinin staining. pSmad1/5+ and BRE:GFP+ cells are primarily localized to cells in the mandibular symphysis and ceratohyal joint that have a rounded, immature morphology and low amounts of ECM secretion. (E-H) Single confocal z-slices of the mandibular symphysis at 72 hpf and 78 hpf from *gdf6a^+/?^*; *Tg(sox10:GFP)* and *gdf6a^-/-^;Tg(sox10:GFP)* animals. Disorganization of the cells in the mandibular symphysis is observed in *gdf6a^-/-^* mutants at 72 hpf and leads to thickening of the symphysis by 78 hpf (arrowhead in H). (I) Quantification of mandibular symphysis length analyzed by unpaired, two-tailed student’s t-test. * = p<0.05. ECM=extracellular matrix, Mc=Meckel’s cartilage, Ch=ceratohyal. Scale bars=50 microns in A, B, C, D, 25 microns in E, G.

### *gdf6a* does not regulate joint marker expression

Gdf6 has previously demonstrated roles in regulating several joint subtypes in the murine skeleton (Settle et al., 2003). Therefore, we reasoned that Gdf6a may be positively regulating joint formation in the ventromedial craniofacial skeleton. We first characterized expression patterns of *nkx3.2,* which is necessary for primary jaw joint development in zebrafish, and *gdf5,* which is expressed at the interzone of the zebrafish primary jaw joint (Miller et al., 2003; S. M. Sperber & Dawid, 2008). Consistent with previous analysis, we observed strong *nkx3.2* expression in the primary jaw joint from 54 hpf until it begins to wane at 72 hpf (Figure S10). Additionally, we observed expression of *nkx3.2* in the midline of the mandibular and hyoid arches in the region of the presumptive mandibular symphysis and ceratohyal joint at 54 hpf, strongest at 60 hpf, and waning at 68 hpf (Figure S10). In contrast to *nkx3.2,* which is highly localized to presumptive joints in the mandibular and hyoid arches, *gdf5* has more widespread expression in the mandibular and hyoid arches (Figure S10). Strong *gdf5* mRNA expression is observed in the perichondrium of the ceratohyal and palatoquadrate from 54 hpf until it begins to fade at 72 hpf (Figure S10). At all stages examined, *gdf5* expression is observed in the ceratohyal joint, but not in the mandibular symphysis (Figure S10). As development proceeds, the expression pattern of *gdf*5 expands in the mandibular arch such that *gdf5* mRNA can be detected in the perichondrium of Meckel’s cartilage (Figure S10). Therefore, we conclude that *nkx3.2* and *gdf5,* two markers of joint formation, are expressed in the zebrafish midline at timepoints relevant to midline morphogenesis and Gdf6a function.

We next examined whether loss of Gdf6a disrupted the expression of joint markers. Compared to siblings, *gdf6a^-/-^* mutants show a mild reduction in *nkx3.2* in the mandibular symphysis and ceratohyal joint at 54 hpf, but the pattern returns to normal by 60 hpf (Figure S10). The expression of *barx1,* a transcription factor that opposes *nkx3.2* function, is unchanged in *gdf6a^-/-^* mutants compared to siblings at 72 hpf (Figure S10) (S. M. Sperber & Dawid, 2008). The expression of *gdf5* is slightly disrupted in the presumptive ceratohyal joint at 54 hpf in *gdf6a^-/-^* mutants (Figure S10), but largely returns to normal by 60 hpf (Figure S10). In summary, Gdf6a has only a modest effect on the expression of joint patterning markers.

### BMP signaling is active in the pharyngeal arches during craniofacial development

In other contexts, GDF6/Gdf6a has been shown to activate canonical BMP signaling via phosphorylation of Smad1/5/8 and transcription of downstream genes regulated by BMP response elements (BREs) (Katagiri et al., 2002). To visualize BMP signaling in the zebrafish pharyngeal arches, we utilized the *Tg(BRE:GFP)* transgenic zebrafish strain (Collery & Link, 2011). Wildtype transgenic animals were fixed at 48, 60, and 72 hpf. At all stages, GFP signal is localized to the midline of the pharyngeal arches in a pattern that corresponds highly to *gdf6a* mRNA expression but appears to peak at 60 hpf (compare Figure 6A-C to Figure 6G-G’’’). Specifically, GFP signal is present in the perioral region, craniofacial cartilage and muscle, and the surrounding mesenchyme (Figure 6A-C). Like *BRE:GFP,* immunofluorescence for pSmad1/5 localizes to the midline of the pharyngeal arches at 48, 60, and 72 hpf in a pattern that closely matches *gdf6a* mRNA expression (Figure 3D-F’’, arrowheads). However, rather than peaking at 60 hpf, pSmad1/5 signal is consistent from 48 to 72 hpf. Additionally, *in-situ* hybridization revealed that the mRNA encoding negative regulators of the BMP signaling pathway, including *smad6a/b* and *bambia/b,* are expressed along the midline of the pharyngeal arches at 60 hpf (Figure S11), Notably, *nog3,* a zebrafish paralog of *NOGGIN/Noggin*, is expressed broadly in the presumptive craniofacial cartilage (Figure 3H-H’’’). In summary, BMP signaling is active in the midline of the pharyngeal arches from 48-72 hpf, with regulators of BMP signaling present in a similar spatiotemporal pattern.

**Figure 6.**
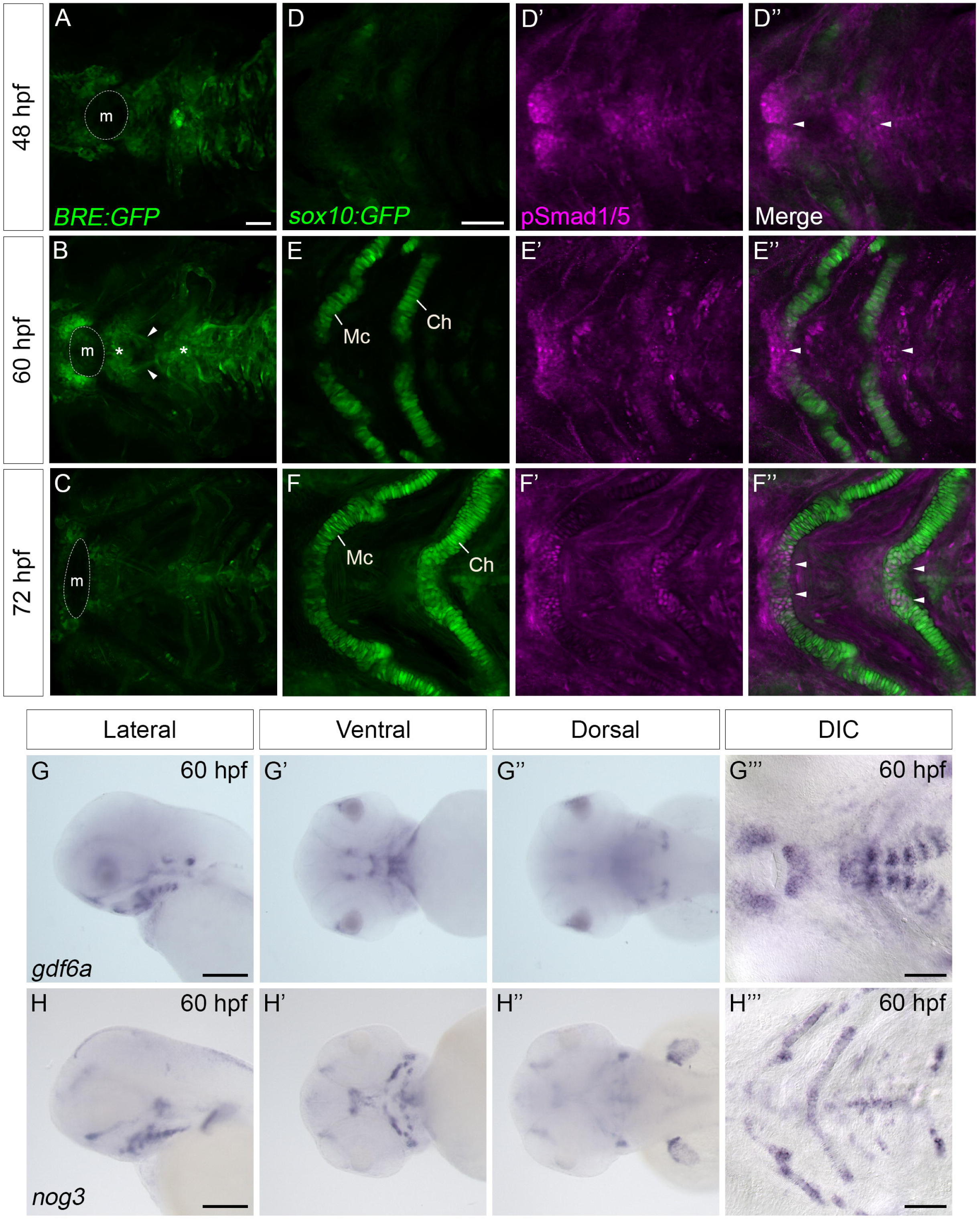
BMP signaling is active in the pharyngeal arches from 48-72 hpf. (A-C) Single confocal z-slices of the pharyngeal arches of *Tg(BRE:GFP)* transgenic zebrafish at 48, 60, and 72 hpf. At 48 hpf, GFP signal is observed in the mesenchyme of the pharyngeal arches and in the perioral region (A). At 60 and 72 hpf, GFP expression is localized to the pharyngeal arch mesenchyme, the craniofacial muscles (arrows), and in the midline of Meckel’s cartilage, the ceratohyal, and the branchial cartilages (asterisks). (D-F’’) Single confocal z-slices of the pharyngeal arches of *Tg(sox10:GFP)* transgenic zebrafish immunolabelled for phosphorylated Smad1/5 (pSmad1/5). pSmad1/5 immunofluorescence recapitulates the pattern of GFP localization observed in *Tg(BRE:GFP)* animals (arrowheads). (G-G’’’) Dissecting microscope (G-G’’) and DIC (G’’’) images of *gdf6a* mRNA expression showing high correlation with *BRE:GFP* transgene activity and pSmad1/5 localization. (H-H’’’) Dissecting microscope (H-H’’) and DIC (H’’’) microscope images of *nog3* mRNA expression in the pharyngeal cartilage at 60 hpf. m=mouth, Mc=Meckel’s cartilage, Ch=ceratohyal. Scale bars=50 microns in A, D, G’’’, H’’’, 100 microns in G, H.

### BMP signaling regulates development of midline structures in the craniofacial skeleton

Thus far, we have established that both *gdf6a* mRNA and readouts of BMP signaling activity are present in the midline of the pharyngeal arches from 48-72 hpf, which correlates with the craniofacial phenotypes of *gdf6a^-/-^* mutant larvae. While the cartilage phenotypes present in *gdf6a^-/-^* mutants are fully penetrant, they are relatively mild compared to other craniofacial phenotypes caused by a loss of signaling ligands, suggesting the potential for compensation from other BMP ligands. Moreover, although *gdf6a* mRNA and BMP activity are localized to the pharyngeal arches from 48-72 hpf, we have not demonstrated the necessity of BMP signaling in formation of midline craniofacial structures during this window. To accomplish this, we utilized the small molecule inhibitor DMH1, which has been shown to inhibit BMP signaling in zebrafish (Hao et al., 2010). We treated wildtype embryos with 10 μM DMH1 or control DMSO-containing media from 48-72 hpf and fixed treated larvae at 5 dpf for Alcian blue and Alizarin red staining. Compared to controls, larvae treated with 10 μM DMH1 exhibited cartilage phenotypes similar to those observed in *gdf6a^-/-^* mutants. At 5 dpf, all DMH1-treated larvae developed misoriented ceratohyals and disorganized mandibular symphyses, indicating that BMP signaling regulates the formation of these structures from 48-72 hpf (Figure 7A-B). Notably, the phenotypes produced by DMH1 treatment were qualitatively more severe compared to the *gdf6a^-/-^* mutant phenotype, suggesting some overlapping activity from other BMP ligands during the development of these structures. When quantified, we observed a statistically significant thickening of the mandibular symphysis, increase in the angle of the ceratohyals, and reduction in the size of the basihyal and hypobranchials in DMH1-treated larvae (Figure 7E, F). Notably, while the hypobranchials are reduced in DMH1-treated larvae compared to controls, the reduction is not as dramatic as the reduction observed between *gdf6a^-/-^* mutants compared to siblings, suggesting that BMP-dependent development of the hypobranchials extends beyond the 48-72 hpf treatment window (Figure 7C-D, G-H). With regards to muscle development, treatment with 10 μM of DMH1 results in aberrant muscle fiber localization similar to *gdf6a^-/-^* mutants without affecting tendon formation, position or morphology (Figure S12). Therefore, we conclude that BMP signaling is necessary for development of the midline structures of the larval craniofacial skeleton, and that multiple ligands likely synergize with Gdf6a to accomplish this.

**Figure 7.**
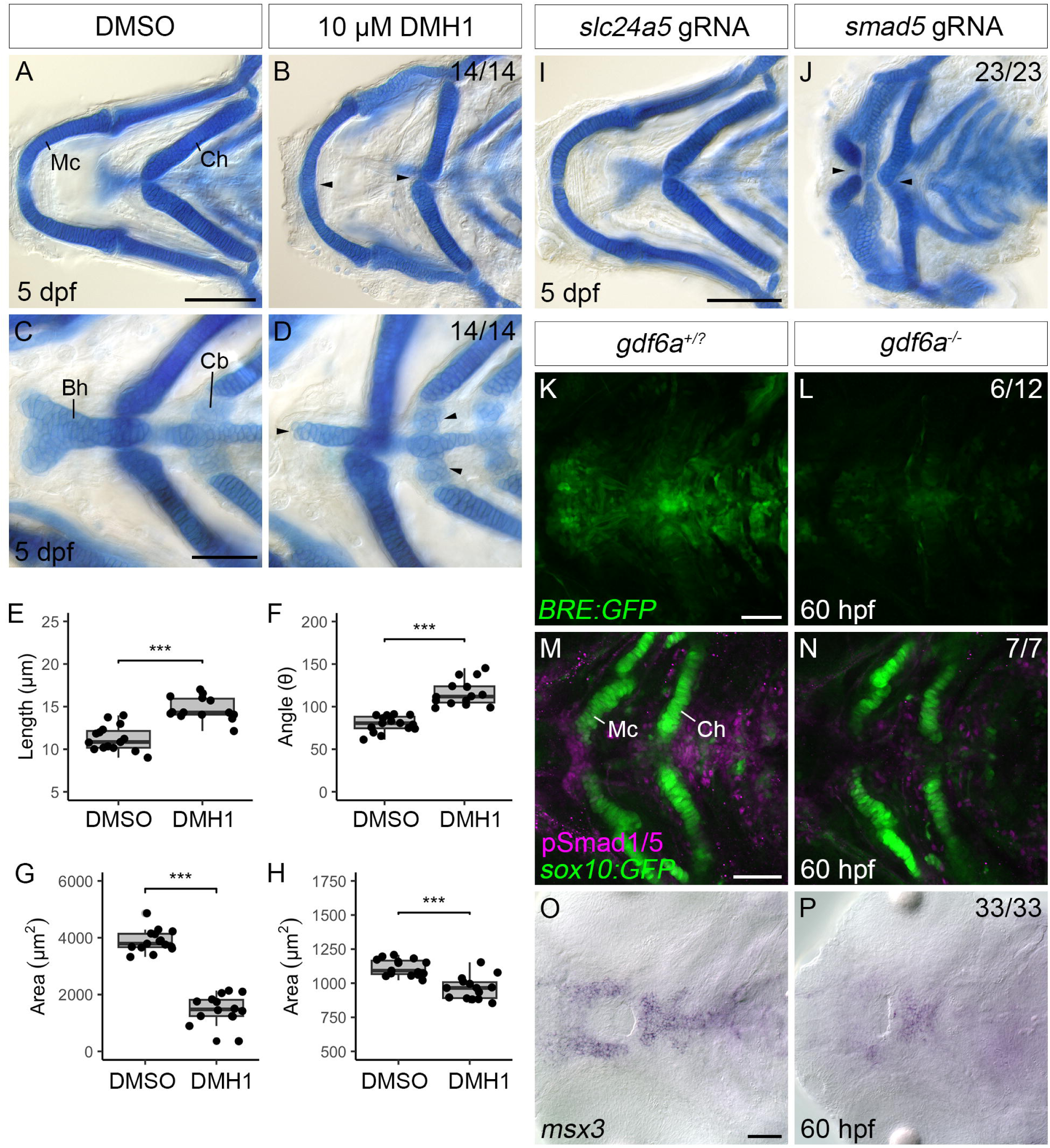
*gdf6a* regulates craniofacial development via BMP signaling. (A-D) DIC microscopy images of flatmounted 5 dpf larvae treated with DMSO or 10μM DMH1 from 48-72 hpf; craniofacial cartilage visualized with Alcian blue and Alizarin red. Larvae treated with DMH1 display several craniofacial abnormalities including thickening of the mandibular symphysis (B, arrowhead), misorientation of the ceratohyals (B, asterisk), malformation of the basihyal (D, arrowhead), and severe reduction of the hypobranchials (D, asterisk). (E-H) Quantification of mandibular symphysis length (E), ceratohyal angle (F), basihyal area (G), and hypobranchial area (H). All analysis used unpaired, two-tailed student’s t-test. Results are representative of N=2 independent experiments. (I-J) Flatmounts of 5 dpf larval craniofacial cartilage visualized with Alcian blue and Alizarin red from larvae injected with *slc24a5* gRNA + Cas9 or *smad5* gRNA + Cas9. *smad5* crispants (J) display defects in the midline of the craniofacial cartilage, including clefting of Meckel’s cartilage (arrow), misorientation of the certaohyals at the midline (arrows), and an absence of the basihyal, basibranchials, and hypobranchials. (K-L) Single confocal z-slices of the pharyngeal arches of *gdf6a^+/?^*;*Tg(BRE:GFP)* and *gdf6a^-/-^*;*Tg(BRE:GFP)* zebrafish. Half (n=6/12) of *gdf6a^-/-^*;*Tg(BRE:GFP)* animals (L) show reduced GFP levels in the pharyngeal arches. (M-N) Single confocal z-slices of the pharyngeal arches of *gdf6a^+/?^*;*(sox10:GFP)* and *gdf6a^-/-^*;*Tg(sox10:GFP)* zebrafish immunolabelled for phosphorylated Smad1/5 (pSmad1/5). (O-P) DIC microscopy images of *msx3* mRNA visualized with *in-situ* hybridization. ***= p<0.001. Mc=Meckel’s cartilage, Ch=ceratohyal, Bh=basihyal, Cb=ceratobranchials. Scale bars=100 microns in A and I, 50 microns in C, K, M, O.

To further test the necessity of BMP signaling in the development of midline craniofacial skeletal structures, we created crispant embryos by injecting wildtype single-cell embryos with a cocktail of Cas9 protein and CRISPR gRNAs targeting either *smad5,* one of the intracellular effectors of BMP signaling, or *slc24a5* as a control. Slc24a5 has a well-known role in pigment production but no known function in craniofacial development (Lamason et al., 2005). Crispant larvae were fixed at 5 dpf and processed for Alcian blue/Alizarin red staining. Compared to *slc24a5* crispants, which showed normal craniofacial anatomy at 5 dpf, all *smad5* crispants displayed an array of craniofacial abnormalities (Figure 7I, J). Specifically, *smad5* crispants displayed clefting of Meckel’s cartilage at the midline and misorientation of the ceratohyals (Figure 7I, J, arrowheads). Furthermore, we also observed an almost complete absence of the basihyal and hypobranchials, indicating that BMP signaling is necessary for the formation of these structures (Figure 7I, J). These phenotypes are consistent with the reported phenotypes of *smad5^-/-^* mutants, demonstrating the specificity and efficiency of our CRISPR/Cas9 protocol (Lovely et al., 2016; Neuhauss et al., 1996; Swartz et al., 2011). Moreover, the Smad5 phenotype is considerably more severe than those present in both *gdf6a^-/-^* mutants and DMH1-treated embryos, further suggesting that multiple BMPs play a role in development of the midline of the zebrafish craniofacial skeleton, especially for the development of the hypobranchials. Taken together, this data indicates that BMP signaling is necessary for the formation and development of the midline of the zebrafish craniofacial skeleton.

### *gdf6a* regulates craniofacial development via BMP signaling

To determine whether Gdf6a regulates pharyngeal arch BMP signaling, we analyzed multiple readouts of BMP signaling in *gdf6a^-/-^* mutants and siblings. Compared to siblings, *gdf6a^-/-^* mutants at 60 hpf show midline reductions in the *BRE:GFP* transgene signal, pSmad 1/5 immunofluorescence, and expression of *msx3,* a transcriptional target of BMP signaling (Figure 7K-P). For all three assessments, the signal was reduced but not completely lost.

The observations that *gdf6a-/-* mutants have only a partial reduction in BMP signaling and weaker phenotypes than larvae with broad BMP inhibition suggest that there is at least some compensatory action from other BMP ligands during development of the midline of the craniofacial skeleton in the mutants. We therefore hypothesized that dampening of BMP signaling would increase the severity of the *gdf6a^-/-^* craniofacial phenotype. We treated *gdf6a^-/-^* mutants and *gdf6a^+/?^* siblings with 5 μM DMH1 (half the concentration used in Figure 7) or DMSO-containing control media from 48-72 hpf, fixed the fish at 7 dpf, and performed Alcian blue and Alizarin red staining. Treatment with 5 μM DMH1 produces a mild, *gdf6a*-like phenotype in most *gdf6a^+/?^* siblings relative to controls, likely owing to the reduction of BMP to sub-threshold in heterozygotes (Figure S13). Furthermore, treatment of *gdf6a^-/-^* mutants with 5 μM DMH1 enhances the *gdf6a^-/-^* mutant phenotype (Figure S13). Qualitatively, this enhancement is observed in severity of the cartilage phenotype; Meckel’s cartilage is fused, the ceratohyals are inverted, and the basihyal and hypobranchials are almost completely absent in DMH1-treated *gdf6a^-/-^* mutants (Figure S13). We also quantified the proportion of embryos that were unaffected, mildly affected, or severely affected and observed that all *gdf6a^-/-^* mutants treated with 5 μM DMH1 presented with the severe phenotype (Figure S13). In contrast, siblings treated with 5 μM DMH1 and DMSO-treated *gdf6a^-/-^* mutants were either unaffected or only mild affected, and siblings treated with DMSO were all unaffected (Figure S13). We observed a similar effect when we performed the same experiment and analyzed muscle fiber localization (Figure S6E-I), suggesting that Gdf6a also synergizes with other sources of BMP signaling during development of the craniofacial muscles. Taken together, this data indicates that other sources of BMP signaling coordinate with Gdf6a during development of midline craniofacial structures.

## Discussion

Although craniofacial abnormalities have been documented in patients with *GDF6* mutations, the underlying etiology remains unclear (Clarke et al., 1994, 1995, 1998; Tassabehji et al., 2008). Here, we demonstrate that Gdf6a, the zebrafish homolog of GDF6, is required for proper formation of midline structures in the craniofacial skeleton.

Using whole-mount *in-situ* hybridization, we observed a striking pattern of *gdf6a* in the pharyngeal arches, with expression along the midline of the craniofacial skeleton in regions that correspond to the developing mandibular symphysis, ceratohyal joint, basibranchial and hypobranchials. Our results are consistent with a previous zebrafish study of transgene expression driven by a pharyngeal arch-specific *gdf6a* promoter (Reed & Mortlock, 2010). The presence and activity of this regulatory element is conserved in mammals, suggesting that the expression pattern of *gdf6a/Gdf6/GDF6* in the pharyngeal arches is highly conserved (Portnoy et al., 2005). This, in turn, suggests Gdf6a has a conserved role in regulating formation of the craniofacial midline despite the divergence in morphology between teleost and mammalian craniofacial skeletons.

Consistent with the expression pattern of *gdf6a* along the ventral midline in the pharyngeal arches, *gdf6a^-/-^* mutants display aberrancies in ventromedial skeletal structures. Specifically, we observe a misorientation of the ceratohyals and an almost complete absence of the hypobranchials, phenotypes previously observed in zebrafish *gdf6a^-/-^* mutants (Reed & Mortlock, 2010). Further, we describe a reduction in the basihyal and disorganization of cells in the mandibular symphysis.

Notably, the loss of hypobranchials in *gdf6a^-/-^* mutants mimics the laryngeal cartilage abnormalities present in patients lacking functional *GDF6*. Indeed, patients with inactivating (but not activating) mutations in *GDF6* present with laryngeal abnormalities, suggesting a necessity for *GDF6* function in laryngeal cartilage formation (Clarke et al., 1994, 1995, 1998). Surprisingly, we found that the loss of laryngeal cartilage was not due to a decrease in cell proliferation or an increase in apoptosis, but resulted from a lack of *sox9a/* Sox9a+ cells in the region of the presumptive hypobranchials, followed by a decrease in *col2a1a* mRNA expression and Col2a1 protein secretion, suggesting that Gdf6a promotes the formation of chondrogenic condensations in the region of the presumptive hypobranchials, and that a lack of chondrogenesis underlies the laryngeal cartilage abnormalities present in patients with a loss of *GDF6*. Similarly, Gdf6a drives chondrogenesis of a cartilage “bridge” connecting the ceratohyals at the midline; in *gdf6a^-/-^* mutants, the bridge is absent and the ceratohyals become misoriented. Our results align with previous studies demonstrating GDF6/Gdf6 promotes chondrogenesis both *in-vitro* and *in-vivo* (Nochi et al., 2004; Shen et al., 2009; Wang et al., 2016; Yu et al., 2023). However, impaired neural crest cell migration could also contribute to loss of the hypobranchials. Indeed, studies in zebrafish and *Xenopus* have shown that *Gdf6/gdf6a* is expressed at the neural plate border and regulates the activity of migratory neural crest cells (Gramann et al., 2019b; Rissi et al., 1995; Schille et al., 2016). We do not favor this explanation, as *gdf6a* is expressed in the pharyngeal arches at positions corresponding to the presumptive hypobranchials starting at 48 hpf, a stage where neural crest migration into the pharyngeal arches has largely ended, but live imaging studies should be done to further investigate the possibility.

Gdf6a regulates mandibular symphysis morphogenesis independent of Sox9a and Col2a1a expression, which contrasts with its function promoting chondrogenesis in pharyngeal arches 2 and 3. We hypothesized that Gdf6a influences PCP in the mandibular symphysis, and its loss disrupts coordinated cell movements, resulting in a thickened symphysis. However, localization of the microtubule organizing center was normal, suggesting PCP is largely intact (Dranow et al., 2023; Le Pabic et al., 2014). While this mechanism cannot be fully excluded, as PCP and non-canonical Wnt signaling can affect Rho GTPases and actin dynamics independent of organelle positioning (Marlow et al., 2002), an alternative explanation is that Gdf6a regulates ECM synthesis or secretion in the symphysis, disrupting chondrocyte morphogenesis. Indeed, proper ECM regulation is known to be critical for craniofacial morphogenesis in zebrafish (Clément et al., 2008; Melville et al., 2011; Sarmah et al., 2010). Further investigation into the cellular role of Gdf6a in mandibular symphysis morphogenesis is therefore warranted.

An analysis of the neurocranium of *gdf6a^-/-^* fish also revealed a role for Gdf6a in development of the ethmoid plate and trabeculae, which are homologous to the mammalian palate. We detected *gdf6a* expression in the region that gives rise to the trabeculae, and the trabeculae in *gdf6a^-/-^* mutants are malformed such that they become detached or almost detached from the rest of the neurocranium. Notably, loss of BMP signaling in the form of either *smad5^-/-^* mutants or an inducible dominant negative type I BMP receptor also causes detachment of the trabeculae (Alexander et al., 2011; Swartz et al., 2011). Moreover, enhancement of BMP signaling results in the overgrowth of cartilage in the trabeculae, indicating that BMP induces chondrogenesis in the maxillary domain (Alexander et al., 2011). Taken together, our evidence implicates Gdf6a as a regulator of trabeculae morphogenesis, which is consistent with previous literature reporting the presence of orofacial clefts and abnormal facial morphology in patients with *GDF6* mutations (Langevin et al., 2014).

We also detected a partially penetrant phenotype of the midline craniofacial muscles in *gdf6a^-/-^* mutants.. Notably, the mislocalization of muscle fibers in *gdf6a^-/-^* mutants occurs independently of alterations to tendons, as tendon structure and marker expression are normal in *gdf6a^-/-^* mutants. While this is at odds with previous studies showing that *GDF6/Gdf6* regulates tendon formation (Mikic et al., 2009; Wolfman et al., 1997), our analysis of BMP signaling using pSmad1/5 immunofluorescence and *BRE:GFP* transgene localization showed BMP activation directly in the muscle fibers, indicating that Gdf6a likely exerts its effect directly on the muscle fibers. While somewhat unexpected, a previous study showed that GDF6 can induce *in-vitro* differentiation of C3H10T1/2 cells into a cardiomyocyte-like fate, suggesting that GDF6 is a potential regulator of muscle formation (Chen et al., 2015).

While we provide evidence that Gdf6a regulates zebrafish craniofacial development via canonical BMP signaling in several structures, including the mandibular symphysis, the ceratohyal joint, and the hypobranchials, our evidence also suggests that Gdf6a is likely working synergistically with other ligands during craniofacial development. This is supported through four lines of evidence: 1) While completely penetrant, the phenotypes present in *gdf6a^-/-^* mutants are relatively subtle, 2) completely inhibiting BMP signaling using DMH1 and *smad5* gRNA + Cas9 produces a more severe phenotype than *gdf6a^-/-^* mutants, 3) there is an incomplete loss of *BRE:GFP* transgene activity and Smad1/5 phosphorylation in *gdf6a^-/-^* mutants, and 4) *gdf6a* mutations synergize with a suboptimal dose of BMP inhibitor to produce more severe craniofacial phenotypes. As is the case with palatal closure, there is evidence from murine models to suggest that other BMP ligands (namely *Bmp2* and *Bmp4*) regulate structures homologous to those affected in *gdf6a^-/-^* mutants, indicating that multiple BMP signals are necessary for craniofacial skeletal development. For instance, mesenchymal deletion of *Bmp4* results in aberrant laryngeal cartilage formation in mouse, including malformation of the thyroid cartilage(Bottasso-Arias et al., 2023). Further investigation is needed to identify the additional BMP ligands that facilitate development of the ventromedial craniofacial skeleton, and their subfunctionality.

Taken together, our research provides an in depth analysis of Gdf6a function during zebrafish craniofacial development and has provided a mechanism by which Gdf6a facilitates the formation of several craniofacial structures frequently malformed in patients with *GDF6* mutations. Future work will focus on elucidating the function of Gdf6a in regulating development of the craniofacial muscles and the trabeculae/ethmoid plate, and whether malformations of these tissues are present in patients with mutations in *GDF6.* This research, in turn, will aid in providing a more complete understanding of Gdf6a’s role in craniofacial development and the pathogenesis of *GDF6-* associated craniofacial abnormalities, thus aiding in the diagnosis, counselling, and management of patients.

## Materials and Methods

### Ethics Statement

Embryonic, larval, and adult zebrafish were cared for according to guidelines set by the Canadian Council of Animal Care (CACC) and protocols were approved by the University of Alberta’s Animal Care and Use Committee (Protocol #427)

### Zebrafish lines and care

Adult zebrafish were kept on a 10-hour dark/14-hour light schedule. Embryos were raised at 25.5°C, 28.5°C or 33°C in embryo media and staged according to standardized developmental hallmarks. Embryos that were grown past 24 hpf were treated with 0.004% 1-phenyl 2-thiourea (PTU) prior to 22 hpf to block pigment formation. Anesthesia of larvae and adult zebrafish was performed with a 4% dilution of 0.4% tricaine methanesulfonate (TMS) stock solution(Prince et al., 1998). The AB wildtype strain, the *gdf6a^s327^* mutant line (Muto et al., 2005), the *Tg(−4.7sox10:GFP)^ba4Tg^* transgenic strain (Dutton et al., 2008b), and the *Tg(5XBMPRE-Xia.Id3:GFP)^ir1189Tg^* transgenic strain (Zuniga et al., 2011) were used in this study.

### Amino acid sequence alignments

Multiple sequence alignment was performed using Clustal Omega (https://www.ebi.ac.uk/jdispatcher/msa/clustalo) and visualized using Jalview software (https://www.jalview.org/). The following polypeptide sequences were retrieved from UniProt for multiple sequence alignment: Gdf5: F1QK08, Gdf6a: P85857, Gdf6b: A8E7N9, Gdf7: A0A8N7TFB7.

### Whole-mount *in-situ* hybridization

Digoxigenin (DIG)-labeled riboprobes were generated from a gene-specific PCR product with an integrated T7 RNA polymerase promoter. The primers used to generate these templates are listed in Table S1. Probe synthesis was performed as previously described (Thisse & Thisse, 2008).

Embryos were fixed in 4% paraformaldehyde (PFA) overnight at 4°C and subsequently permeabilized by incubating embryos in ice-cold 100% acetone for 20 minutes at −20°C. *In-situ* hybridization was performed essentially as previously described except probes were not hydrolyzed (Prince et al., 1998). After coloration was complete, the coloration reaction was stopped by incubating embryos in 100% methanol overnight at 4°C. Embryos were then rehydrated into phosphate buffered saline + 0.1% Tween-20 (PBST) for imaging. For whole-mount imaging, embryos were mounted in 3% methylcellulose and imaged using a Leica MZ 16 F stereomicroscope with a Zeiss Axiocam 305 color digital camera. For DIC imaging, embryos were passed through a series of 30%, 50%, and 70% glycerol, mounted under a coverslip in 70% glycerol, and imaged on a Zeiss AxioImager Z1 compound microscope with a Zeiss Axiocam HR digital camera.

### Alcian blue and Alizarin red histology

Alcian blue and Alizarin red histology was performed as previously described (Walker & Kimmel, 2007). After histology, larvae were transferred to 70% glycerol. For whole-mount imaging, larvae were transferred to a spot plate in 70% glycerol and imaged using a Leica MZ 16 F stereomicroscope with a Zeiss Axiocam 305 color digital camera. For DIC imaging, the skeletal elements of interest were dissected, mounted under a coverslip in 70% glycerol, and imaged using a Zeiss AxioImager Z1 compound microscope with an Axiocam HR digital camera.

### Phalloidin staining

For phalloidin staining, embryos were fixed in 4% PFA overnight at 4°C. Larvae were then washed in PBST before being washed twice in Tris-buffered saline + 0.1% Tween 20 (TBST). Larvae were then permeabilized in 4% Triton X-100 in TBST for 1 hour at room temperature. Larvae were then washed and incubated in Alexa Fluor 488 Phalloidin diluted in TBST (1:100, Life Technologies A12379) overnight at 4°C. After phalloidin staining, larvae were passed through a series of 30%, 50%, and 70% glycerol. Larvae were then mounted on glass slides under a coverslip in 70% glycerol and imaged on a Zeiss AxioImager Z1 compound microscope using confocal imaging.

### Wheat germ agglutinin staining

For wheat germ agglutinin staining, embryos were fixed in 4% PFA overnight at 4°C. Larvae were then washed in phosphate-buffered saline + 0.15% Triton X-100 (PBSTX) and incubated in Alexa-Fluor 647-conjugated wheat germ agglutinin diluted in BPSTX (1:100, Invitrogen W32466) overnight at 4°C. Embryos were then washed in PBSTX and passed through a series of 30%, 50%, and 70% glycerol. Embryos were then mounted on glass slides under a coverslip in 70% glycerol and imaged on a Zeiss AxioImager Z1 compound microscope using confocal imaging.

### Immunofluorescence

For phosphorylated Smad1/5 (pSmad1/5) and Sox9a immunofluorescence, embryos were fixed in 4% PFA overnight at 4°C. pSmad1/5 and Sox9a immunofluorescence was then performed as previously described, except that embryos were then permeabilized in ice-cold acetone for 20 minutes at −20°C instead of using proteinase K (French et al., 2009). After permeabilization, embryos were blocked for at least 1 hour at room temperature using 10% goat serum and 1% bovine serum albumin (BSA) in PBST. Embryos were incubated in rabbit anti-pSmad1/5 (1:200, Cell Signaling Technologies 9511) or mouse anti-Sox9 (1:200, Abcam ab76997) primary antibodies diluted in block overnight at 4°C. After washing, embryos were incubated in goat anti-rabbit Alexa Fluor 647 Plus (1:1000, Life Technologies A32733) or goat anti-mouse Alexa Fluor 488 Plus (1:1000, Life Technologies A32723) secondary antibody overnight at 4°C.

For Thrombospondin 4 (Tbs4) immunofluorescence, embryos were fixed in 95% methanol/5% glacial acetic acid for 4-6 hours at room temperature. Tbs4 immunofluorescence was performed essentially as previously described (Subramanian & Schilling, 2014). After fixation, embryos were permeabilized in ice-cold acetone for 20 minutes at −20°C. Embryos were then blocked for at least 1 hour at room temperature using 5% goat serum and 1% BSA in 0.5% PBSDT. Embryos were incubated in rabbit anti-Tbs4 primary antibody diluted in block (1:400, Abcam ab211143) overnight at 4°C. After washing, embryos were incubated in goat anti-rabbit Alexa Fluor 488 Plus antibody (1:1000, Life Technologies A32731) overnight at 4°C.

For Col2a1a immunofluorescence, embryos were fixed in 4% PFA overnight at 4°C. Col2a1a immunofluorescence was then performed as previously described, except embryos were then washed and permeabilized in ice-cold acetone for 20 minutes at −20°C. Embryos were then blocked for at least 1 hour at room temperature in 2% goat serum and 2 mg/ml BSA in PBST. Embryos were incubated in rabbit anti-Col2a1 primary antibody diluted in block (1:200, Abcam ab209865) overnight at 4°C. After washing, embryos were incubated in goat anti-rabbit Alexa Fluor 488 Plus antibody (1:1000, Life Technologies A32731) overnight at 4°C.

For phosphorylated Histone H3 (pH3) immunofluorescence, embryos were fixed in 4% PFA overnight at 4°C. Embryos were then washed and permeabilized in ice-cold acetone for 20 minutes at −20°C. After permeabilization, embryos were washed, and antigen retrieval was performed by incubating embryos in 10 mM citric acid at 95°C for 10 minutes. Embryos were then blocked for at least 1 hour using 3% BSA in phosphate buffered saline with 0.5% Triton X-100 (0.5% PBSTX). Embryos were incubated in rabbit anti-pH3 primary antibody diluted in block (1:1000, Abcam ab32107) for 2 hours at room temperature. After washing, embryos were incubated in goat anti-rabbit Alexa Fluor 488 Plus antibody (1:1000, Life Technologies A32731) and TO-PRO-3 Iodide (642/661) (1:2000, Invitrogen T3605) for 2 hours at room temperature.

For cleaved Caspase3 immunofluorescence, embryos were fixed in 4% PFA overnight at 4°C. Embryos were then washed and permeabilized in ice-cold acetone for 20 minutes at −20°C. Embryos were washed and blocked for at least 30 minutes in 5% goat serum in PBSDTT. Embryos were incubated in rabbit anti-cleaved Caspase3 primary antibody (1:400, BD Pharminigen 559565). After washing, embryos were incubated in goat anti-rabbit Alexa Fluor 488 Plus antibody (1:1000, Life Technologies A32731) and TO-PRO-3 Iodide (642/661) (1:2000, Invitrogen T3605) for 2 hours at room temperature.

After immunofluorescence, embryos were passed through a series of 30%, 50%, and 70% glycerol. Embryos were then mounted on glass slides under a coverslip in 70% glycerol and imaged on a Zeiss AxioImager Z1 compound microscope using confocal imaging.

### Pharmacological treatments

Prior to treatment, embryos were dechorionated and treated with PTU to prevent pigment formation (if applicable). At 48 hpf, embryos were transferred to 35 mm petri dishes (n≤15) and treated with either 5 or 10 μM of DMH1 (Millipore Sigma, D8946) or an equivalent volume of DMSO dissolved in embryo media. Embryos were then incubated at 28.5°C for the duration of treatment. At 72 hpf, embryo media containing DMH1 or DMSO was removed, and embryos were washed three times with fresh zebrafish embryo media. Embryos were then returned to 28.5°C until fixed for downstream assays.

### G0 CRISPR analysis

Gene-specific templates for guide RNA (gRNA) were taken from a previously published list of sequences for zebrafish genes and are listed in Table S2. gRNA synthesis was performed essentially as previously described (Wu et al., 2018). Briefly, gene-specific gRNA template oligos were annealed to a core oligo, filled in using T4 DNA polymerase (Invitrogen EP0061), and PCR purified. The annealed gRNA templates were then used for pooled gRNA synthesis with the MEGAscript T7 Transcription Kit (Ambion AM1334). RNA was then precipitated and resuspended in RNAse-free water and the concentration was determined using a spectrophotometer. gRNA was then diluted to a final concentration of 0.5 μg/μL and stored a-80°C until injection.

Prior to injection, gRNA was thawed on ice. After thawing, 0.5 μL TrueCut Cas9 Protein v2 (Invitrogen A36498) was added to the gRNA mixture. 2 nL of Cas9/gRNA mixture was injected directly into the cell of single-cell embryos. After injection, embryos were transferred to embryo media and used for downstream assays.

### Data analysis and statistics

ImageJ software was used for all measurements (https://imagej.net/ij/). For anatomical measurements, the “Straight” tool was used to measure length, the “Angle” tool was used to measure angles, and the “Polygon selections” tool was used to measure area. Quantification of γ-Tubulin localization was performed as previously described (Dranow et al., 2023). All data was recorded in Microsoft Excel.

Statistical tests were conducted in Microsoft Excel. The following statistical tests were used: Student’s t-test, Chi square test. All graphs were made using the ggplot2 package in R or Microsoft Excel.

## Acknowledgements

The authors thank the Molecular Biology Service Unit (MBSU) and Science Animal Support Services (SASS) at the University of Alberta for use of core facilities and animal care, respectively.

## Data and resource availability

All relevant data and details of resources can be found within the article and its supplementary information.

## References

Alexander, C., Zuniga, E., Blitz, I. L., Wada, N., Le Pabic, P., Javidan, Y., Zhang, T., Cho, K. W., Crump, J. G., & Schilling, T. F. (2011). Combinatorial roles for BMPs and Endothelin 1 in patterning the dorsal-ventral axis of the craniofacial skeleton. Development, 138(23), 5135–5146. 10.1242/dev.067801

Bottasso-Arias, N., Burra, K., Sinner, D., & Riede, T. (2023). Disruption of BMP4 signaling is associated with laryngeal birth defects in a mouse model. Developmental Biology, 500, 10– 21. 10.1016/J.YDBIO.2023.04.007

Brunet, L. J., McMahon, J. A., McMahon, A. P., & Harland, R. M. (1998). Noggin, Cartilage Morphogenesis, and Joint Formation in the Mammalian Skeleton. Science, 280(5368), 1455–1457. 10.1126/science.280.5368.1455

Chang, S. C., Hoang, B., Thomas, J. T., Vukicevic, S., Luyten, F. P., Ryba, N. J., Kozak, C. A., Reddi, A. H., & Moos, M. (1994). Cartilage-derived morphogenetic proteins. New members of the transforming growth factor-beta superfamily predominantly expressed in long bones during human embryonic development. Journal of Biological Chemistry, 269(45), 28227–28234. 10.1016/S0021-9258(18)46918-9

Chen, L., Wei, H., Tan, J., Chen, H., Liu, Z., & Chen, Y. (2015). Bone morphogenetic protein 9 and 13 induce C3H10T1/2 cell differentiation to cardiomyocyte-like cells in vitro. Cell Transplantation, 24(5), 909–920. 10.3727/096368913X676907/ASSET/IMAGES/LARGE/10.3727_096368913X676907-FIG6.JPEG

Clarke, R. A., Catalan, G., Diwan, A. D., & Kearsley, J. H. (1998). Heterogeneity in Klippel-Feil syndrome: A new classification. Pediatric Radiology, 28(12), 967–974. 10.1007/S002470050511/METRICS

Clarke, R. A., Davis, P. J., & Tonkin, J. (1994). Klippel-Feil Syndrome Associated with Malformed Larynx: Case Report. Annals of Otology, Rhinology & Laryngology, 103(3), 201–207. 10.1177/000348949410300306

Clarke, R. A., Singh, S., McKenzie, H., Kearsley, J. H., & Yip, M. Y. (1995). Familial Klippel-Feil syndrome and paracentric inversion inv(8)(q22.2q23.3). American Journal of Human Genetics, 57(6), 1364. https://pmc.ncbi.nlm.nih.gov/articles/PMC1801422/

Clément, A., Wiweger, M., von der Hardt, S., Rusch, M. A., Selleck, S. B., Chien, C.-B., & Roehl, H. H. (2008). Regulation of Zebrafish Skeletogenesis by ext2/dackel and papst1/pinscher. PLOS Genetics, 4(7), e1000136. 10.1371/journal.pgen.1000136

Clendenning, D. E., & Mortlock, D. P. (2012). The BMP Ligand Gdf6 Prevents Differentiation of Coronal Suture Mesenchyme in Early Cranial Development. PLOS ONE, 7(5), e36789. 10.1371/JOURNAL.PONE.0036789

Collery, R. F., & Link, B. A. (2011). Dynamic Smad-mediated BMP signaling revealed through transgenic zebrafish. Developmental Dynamics : An Official Publication of the American Association of Anatomists, 240(3), 712. 10.1002/DVDY.22567

Cubbage, C. C., & Mabee, P. M. (1996). Development of the cranium and paired fins in the zebrafish Danio rerio (Ostariophysi, Cyprinidae). Journal of Morphology, 229(2), 121–160. 10.1002/(SICI)1097-4687(199608)229:2<121::AID-JMOR1>3.0.CO;2-4

Dranow, D. B., Le Pabic, P., & Schilling, T. F. (2023). The non-canonical Wnt receptor Ror2 is required for cartilage cell polarity and morphogenesis of the craniofacial skeleton in zebrafish. Development, 150(8), dev201273. 10.1242/dev.201273

Ducy, P., & Karsenty, G. (2000). The family of bone morphogenetic proteins. Kidney International, 57(6), 2207–2214. 10.1046/J.1523-1755.2000.00081.X

Dutton, J. R., Antonellis, A., Carney, T. J., Rodrigues, F. S. L. M., Pavan, W. J., Ward, A., & Kelsh, R. N. (2008a). An evolutionarily conserved intronic region controls the spatiotemporal expression of the transcription factor Sox10. BMC Developmental Biology, 8(1), 105. 10.1186/1471-213X-8-105

Dutton, J. R., Antonellis, A., Carney, T. J., Rodrigues, F. S. L. M., Pavan, W. J., Ward, A., & Kelsh, R. N. (2008b). An evolutionarily conserved intronic region controls the spatiotemporal expression of the transcription factor Sox10. BMC Developmental Biology, 8(1), 1–20. 10.1186/1471-213X-8-105/TABLES/5

French, C. R., Erickson, T., French, D. V, Pilgrim, D. B., & Waskiewicz, A. J. (2009). Gdf6a is required for the initiation of dorsal–ventral retinal patterning and lens development. Developmental Biology, 333(1), 37–47.

Gorlin, R. J., Cohen Jr, M. M., & Hennekam, R. C. M. (2001). Syndromes of the head and neck. Oxford university press.

Gosse, N. J., & Baier, H. (2009). An essential role for Radar (Gdf6a) in inducing dorsal fate in the zebrafish retina. Proceedings of the National Academy of Sciences, 106(7), 2236 LP – 2241. 10.1073/pnas.0803202106

Gramann, A. K., Venkatesan, A. M., Guerin, M., & Ceol, C. J. (2019). Regulation of zebrafish melanocyte development by ligand-dependent BMP signaling. ELife, 8, e50047. 10.7554/eLife.50047

Groppe, J., Greenwald, J., Wiater, E., Rodriguez-Leon, J., Economides, A. N., Kwiatkowski, W., Affolter, M., Vale, W. W., Belmonte, J. C. I., & Choe, S. (2002). Structural basis of BMP signalling inhibition by the cystine knot protein Noggin. Nature, 420(6916), 636–642. 10.1038/nature01245

Hao, J., Ho, J. N., Lewis, J. A., Karim, K. A., Daniels, R. N., Gentry, P. R., Hopkins, C. R., Lindsley, C. W., & Hong, C. C. (2010). In vivo structure - Activity relationship study of dorsomorphin analogues identifies selective VEGF and BMP inhibitors. ACS Chemical Biology, 5(2), 245–253. 10.1021/CB9002865/SUPPL_FILE/CB9002865_SI_007.PDF

Imamura, T., Takase, M., Nishihara, A., Oeda, E., Hanai, J., Kawabata, M., & Miyazono, K. (1997). Smad6 inhibits signalling by the TGF-β superfamily. Nature, 389(6651), 622–626. 10.1038/39355

Jiang, X., Iseki, S., Maxson, R. E., Sucov, H. M., & Morriss-Kay, G. M. (2002). Tissue origins and interactions in the mammalian skull vault. Developmental Biology, 241(1), 106–116.

Katagiri, T., Imada, M., Yanai, T., Suda, T., Takahashi, N., & Kamijo, R. (2002). Identification of a BMP-responsive element in Id1, the gene for inhibition of myogenesis. Genes to Cells, 7(9), 949–960. 10.1046/j.1365-2443.2002.00573.x

Kawabata, M., Inoue, H., Hanyu, A., Imamura, T., & Miyazono, K. (1998). Smad proteins exist as monomers in vivo and undergo homo- and hetero-oligomerization upon activation by serine/threonine kinase receptors. The EMBO Journal, 17(14), 4056–4065. 10.1093/emboj/17.14.4056

Lamason, R. L., Mohideen, M. A. P. K., Mest, J. R., Wong, A. C., Norton, H. L., Aros, M. C., Jurynec, M. J., Mao, X., Humphreville, V. R., Humbert, J. E., Sinha, S., Moore, J. L., Jagadeeswaran, P., Zhao, W., Ning, G., Makalowska, I., McKeigue, P. M., O’Donnell, D., Kittles, R.,… Cheng, K. C. (2005). Genetics: SLC24A5, a putative cation exchanger, affects pigmentation in zebrafish and humans. Science, 310(5755), 1782–1786. 10.1126/SCIENCE.1116238

Langevin, K. M., Mabry, K., & Castiglione, C. (2014). An uncommon case: Cleft palate, respiratory compromise, and klippel-feil anomaly. Plastic Surgical Nursing, 34(3), 127– 132. 10.1097/PSN.0000000000000052

Le Pabic, P., Ng, C., & Schilling, T. F. (2014). Fat-Dachsous Signaling Coordinates Cartilage Differentiation and Polarity during Craniofacial Development. PLOS Genetics, 10(10), e1004726. 10.1371/journal.pgen.1004726

Lièvre, C. L., & Douarin, N. L. (1975). Mesenchymal derivatives of the neural crest: analysis of chimaeric quail and chick embryos. Development, 34(1), 125–154.

Lovely, C. Ben, Swartz, M. E., McCarthy, N., Norrie, J. L., & Eberhart, J. K. (2016). Bmp signaling mediates endoderm pouch morphogenesis by regulating Fgf signaling in zebrafish. Development, 143(11), 2000–2011. 10.1242/dev.129379

Marlow, F., Topczewski, J., Sepich, D., & Solnica-Krezel, L. (2002). Zebrafish Rho kinase 2 acts downstream of Wnt11 to mediate cell polarity and effective convergence and extension movements. Current Biology, 12(11), 876–884. 10.1016/S0960-9822(02)00864-3

McGaughran, J. M., Kuna, P., & Das, V. (1998). Audiological abnormalities in the Klippel-Feil syndrome. Archives of Disease in Childhood, 79(4), 352–355. 10.1136/ADC.79.4.352

Melville, D. B., Montero-Balaguer, M., Levic, D. S., Bradley, K., Smith, J. R., Hatzopoulos, A. K., & Knapik, E. W. (2011). The feelgood mutation in zebrafish dysregulates COPII-dependent secretion of select extracellular matrix proteins in skeletal morphogenesis. Disease Models & Mechanisms, 4(6), 763–776. 10.1242/dmm.007625

Mikic, B., Rossmeier, K., & Bierwert, L. (2009). Identification of a Tendon Phenotype in GDF6 Deficient Mice. The Anatomical Record: Advances in Integrative Anatomy and Evolutionary Biology, 292(3), 396–400. 10.1002/AR.20852

Miller, C. T., Yelon, D., Stainier, D. Y. R., & Kimmel, C. B. (2003). Two endothelin 1 effectors, hand2 and bapx1, pattern ventral pharyngeal cartilage and the jaw joint.

Miyamoto, Y., Mabuchi, A., Shi, D., Kubo, T., Takatori, Y., Saito, S., Fujioka, M., Sudo, A., Uchida, A., & Yamamoto, S. (2007). A functional polymorphism in the 5′ UTR of GDF5 is associated with susceptibility to osteoarthritis. Nature Genetics, 39(4), 529–533.

Mork, L., & Crump, G. (2015). Chapter Ten - Zebrafish Craniofacial Development: A Window into Early Patterning. In Y. B. T.-C. T. in D. B. Chai (Ed.), Craniofacial Development (Vol. 115, pp. 235–269). Academic Press. 10.1016/bs.ctdb.2015.07.001

Muto, A., Orger, M. B., Wehman, A. M., Smear, M. C., Kay, J. N., Page-McCaw, P. S., Gahtan, E., Xiao, T., Nevin, L. M., Gosse, N. J., Staub, W., Finger-Baier, K., & Baier, H. (2005). Forward Genetic Analysis of Visual Behavior in Zebrafish. PLOS Genetics, 1(5), e66. 10.1371/JOURNAL.PGEN.0010066

Nakao, A., Afrakhte, M., Morn, A., Nakayama, T., Christian, J. L., Heuchel, R., Itoh, S., Kawabata, M., Heldin, N.-E., Heldin, C.-H., & Dijke, P. ten. (1997). Identification of Smad7, a TGFβ-inducible antagonist of TGF-β signalling. Nature, 389(6651), 631–635. 10.1038/39369

Neuhauss, S. C. F., Solnica-Krezel, L., Schier, A. F., Zwartkruis, F., Stemple, D. L., Malicki, J., Abdelilah, S., Stainier, D. Y. R., & Driever, W. (1996). Mutations affecting craniofacial development in zebrafish. Development, 123(1), 357–367. 10.1242/DEV.123.1.357

Nochi, H., Jin, H. S., Lou, J., Adkisson, H. D., Maloney, W. J., & Hruska, K. A. (2004). Adenovirus Mediated BMP 13 Gene Transfer Induces Chondrogenic Differentiation of Murine Mesenchymal Progenitor Cells. Journal of Bone and Mineral Research, 19(1), 111– 122. 10.1359/JBMR.2004.19.1.111

Noden, D. M. (1978). The control of avian cephalic neural crest cytodifferentiation: I. Skeletal and connective tissues. Developmental Biology, 67(2), 296–312.

Onichtchouk, D., Chen, Y.-G., Dosch, R., Gawantka, V., Delius, H., Massagué, J., & Niehrs, C. (1999). Silencing of TGF-β signalling by the pseudoreceptor BAMBI. Nature, 401(6752), 480–485. 10.1038/46794

Portnoy, M. E., McDermott, K. J., Antonellis, A., Margulies, E. H., Prasad, A. B., Kingsley, D. M., Green, E. D., & Mortlock, D. P. (2005). Detection of potential GDF6 regulatory elements by multispecies sequence comparisons and identification of a skeletal joint enhancer. Genomics, 86(3), 295–305. 10.1016/J.YGENO.2005.05.003

Prince, V. E., Moens, C. B., Kimmel, C. B., & Ho, R. K. (1998). Zebrafish hox genes: expression in the hindbrain region of wild-type and mutants of the segmentation gene, valentino. Development, 125(3), 393 LP – 406. http://dev.biologists.org/content/125/3/393.abstract

Reed, N. P., & Mortlock, D. P. (2010). Identification of a distant cis-regulatory element controlling pharyngeal arch-specific expression of zebrafish gdf6a/radar. Developmental Dynamics, 239(4), 1047–1060. 10.1002/dvdy.22251

Rissi, M., Wittbrodt, J., Délot, E., Naegeli, M., & Rosa, F. M. (1995). Zebrafish Radar: A new member of the TGF-β superfamily defines dorsal regions of the neural plate and the embryonic retina. Mechanisms of Development, 49(3), 223–234. 10.1016/0925-4773(94)00320-M

Sarmah, S., Barrallo-Gimeno, A., Melville, D. B., Topczewski, J., Solnica-Krezel, L., & Knapik, E. W. (2010). Sec24D-Dependent Transport of Extracellular Matrix Proteins Is Required for Zebrafish Skeletal Morphogenesis. PLOS ONE, 5(4), e10367. 10.1371/journal.pone.0010367

Schille, C., Bayerlová, M., Bleckmann, A., & Schambony, A. (2016). Ror2 signaling is required for local upregulation of GDF6 and activation of BMP signaling at the neural plate border. Development, 143(17), 3182–3194. 10.1242/DEV.135426

Schilling, T. F., & Kimmel, C. B. (1994). Segment and cell type lineage restrictions during pharyngeal arch development in the zebrafish embryo. Development, 120(3), 483–494. 10.1242/dev.120.3.483

Settle, S. H., Rountree, R. B., Sinha, A., Thacker, A., Higgins, K., & Kingsley, D. M. (2003). Multiple joint and skeletal patterning defects caused by single and double mutations in the mouse Gdf6 and Gdf5 genes. Developmental Biology, 254(1), 116–130. 10.1016/S0012-1606(02)00022-2

Shen, B., Bhargav, D., Wei, A., Williams, L. A., Tao, H., Ma, D. D. F., & Diwan, A. D. (2009). BMP-13 Emerges as a Potential Inhibitor of Bone Formation. International Journal of Biological Sciences, 5(2), 192–200. 10.7150/IJBS.5.192

Sperber, G. H., Sperber, S. M., & Guttmann, G. D. . (2010). Craniofacial embryogenetics and development (2nd ed.). People’s Medical Pub. House USA.

Sperber, S. M., & Dawid, I. B. (2008). barx1 is necessary for ectomesenchyme proliferation and osteochondroprogenitor condensation in the zebrafish pharyngeal arches. Developmental Biology, 321(1), 101–110. 10.1016/J.YDBIO.2008.06.004

Storm, E. E., Huynh, T. V, Copeland, N. G., Jenkins, N. A., Kingsley, D. M., & Lee, S.-J. (1994). Limb alterations in brachypodism mice due to mutations in a new member of the TGFβ-superfamily. Nature, 368(6472), 639–643. 10.1038/368639a0

Subramanian, A., & Schilling, T. F. (2014). Thrombospondin-4 controls matrix assembly during development and repair of myotendinous junctions. ELife, 2014(3). 10.7554/ELIFE.02372

Swartz, M. E., Sheehan-Rooney, K., Dixon, M. J., & Eberhart, J. K. (2011). Examination of a palatogenic gene program in zebrafish. Developmental Dynamics, 240(9), 2204–2220. 10.1002/dvdy.22713

Tassabehji, M., Fang, Z. M., Hilton, E. N., McGaughran, J., Zhao, Z., de Bock, C. E., Howard, E., Malass, M., Donnai, D., Diwan, A., Manson, F. D. C., Murrell, D., & Clarke, R. A. (2008). Mutations in GDF6 are associated with vertebral segmentation defects in Klippel-Feil syndrome. Human Mutation, 29(8), 1017–1027. 10.1002/humu.20741

Thisse, C., & Thisse, B. (2008). High-resolution in situ hybridization to whole-mount zebrafish embryos. Nature Protocols, 3(1), 59–69. 10.1038/nprot.2007.514

Valdes, A. M., Evangelou, E., Kerkhof, H. J. M., Tamm, A., Doherty, S. A., Kisand, K., Tamm, A., Kerna, I., Uitterlinden, A., & Hofman, A. (2011). The GDF5 rs143383 polymorphism is associated with osteoarthritis of the knee with genome-wide statistical significance. Annals of the Rheumatic Diseases, 70(5), 873–875.

Walker, M., & Kimmel, C. (2007). A two-color acid-free cartilage and bone stain for zebrafish larvae. Biotechnic & Histochemistry, 82(1), 23–28. 10.1080/10520290701333558

Wang, J., Yu, T., Wang, Z., Ohte, S., Yao, R., Zheng, Z., Geng, J., Cai, H., Ge, Y., Li, Y., Xu, Y., Zhang, Q., Gusella, J. F., Fu, Q., Pregizer, S., Rosen, V., & Shen, Y. (2016). A New Subtype of Multiple Synostoses Syndrome Is Caused by a Mutation in GDF6 That Decreases Its Sensitivity to Noggin and Enhances Its Potency as a BMP Signal. Journal of Bone and Mineral Research, 31(4), 882–889. 10.1002/jbmr.2761

Warren, S. M., Brunet, L. J., Harland, R. M., Economides, A. N., & Longaker, M. T. (2003). The BMP antagonist noggin regulates cranial suture fusion. Nature, 422(6932), 625–629. 10.1038/nature01545

Wolfman, N. M., Hattersley, G., Cox, K., Celeste, A. J., Nelson, R., Yamaji, N., Dube, J. L., DiBlasio-Smith, E., Nove, J., Song, J. J., Wozney, J. M., & Rosen, V. (1997). Ectopic induction of tendon and ligament in rats by growth and differentiation factors 5, 6, and 7, members of the TGF-beta gene family. The Journal of Clinical Investigation, 100(2), 321– 330. 10.1172/JCI119537

Wu, R. S., Lam, I. I., Clay, H., Duong, D. N., Deo, R. C., & Coughlin, S. R. (2018). A Rapid Method for Directed Gene Knockout for Screening in G0 Zebrafish. Developmental Cell, 46(1), 112–125.e4. 10.1016/j.devcel.2018.06.003

Yamashita, H., ten Dijke, P., Franzen, P., Miyazono, K., & Heldin, C.-H. (1994). Formation of hetero-oligomeric complexes of type I and type II receptors for transforming growth factor-beta. Journal of Biological Chemistry, 269(31), 20172–20178.

Yoshida, T., Vivatbutsiri, P., Morriss-Kay, G., Saga, Y., & Iseki, S. (2008). Cell lineage in mammalian craniofacial mesenchyme. Mechanisms of Development, 125(9), 797–808. 10.1016/j.mod.2008.06.007

Yu, T., Li, G., Wang, C., Li, N., Yao, R., & Wang, J. (2023). Defective Joint Development and Maintenance in GDF6-Related Multiple Synostoses Syndrome. Journal of Bone and Mineral Research, 38(4), 568–577. 10.1002/jbmr.4785

Zimmerman, L. B., De Jesús-Escobar, J. M., & Harland, R. M. (1996). The Spemann organizer signal noggin binds and inactivates bone morphogenetic protein 4. Cell, 86(4), 599–606. 10.1016/S0092-8674(00)80133-6

Zuniga, E., Rippen, M., Alexander, C., Schilling, T. F., & Crump, J. G. (2011). Gremlin 2 regulates distinct roles of BMP and Endothelin 1 signaling in dorsoventral patterning of the facial skeleton. Development, 138(23), 5147–5156. 10.1242/dev.067785

