## Supplementary Information for "The BMP ligand Gdf6a Regulates Development of the Zebrafish Craniofacial Skeleton in a Pharyngeal Arch-Specific Manner"

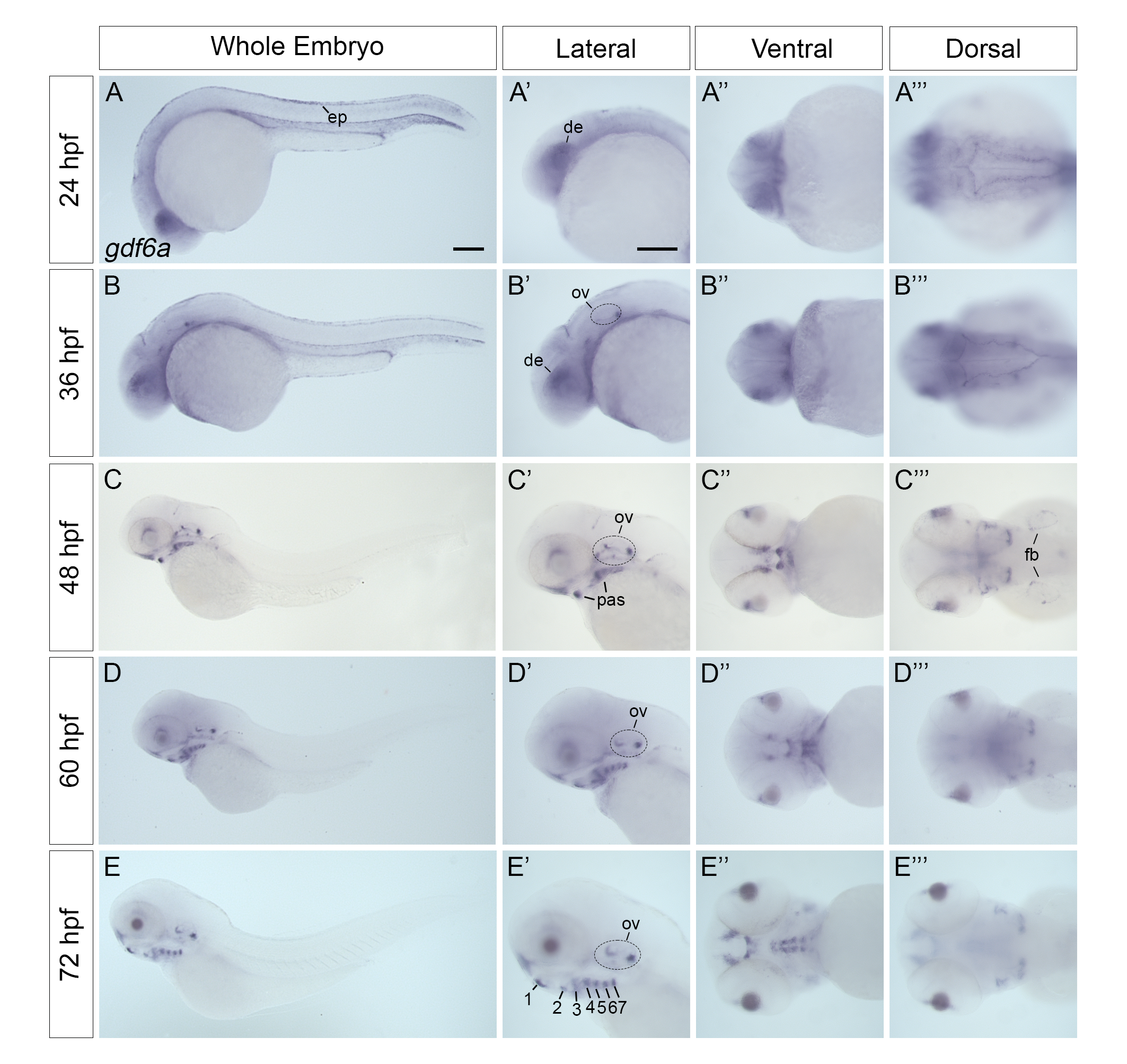
**Figure S1 – *gdf6a* expression pattern in 24-72 hpf zebrafish.** (A-E’’’) *in situ* hybridization showing *gdf6a* mRNA localization in several tissues during zebrafish development. At the early pharyngula stages (24-36 hpf), *gdf6a* is localized to the epidermis and dorsal eye (A-B’’’). At 36 hpf, *gdf6a* is also localized to the ventral head mesenchyme and otic vesicle. As development proceeds, *gdf6a* mRNA expression begins in the pharyngeal arches and perioral region (C-C’’). At this stage, *gdf6a* expression also begins in the fin buds (C’’’) and expression in the otic vesicle intensifies (C, C’, C’’’). The general pattern of *gdf6a* mRNA expression in the pharyngeal arches, perioral region, and otic vesicle is maintained through the late pharyngula stages (60 hpf, D-D’’’) and the early larval stages (72 hpf, E-E’’’). Notably, *gdf6a* is expressed segmentally in each pharyngeal arch (E’). ep=epidermis, de=dorsal eye, ov=otic vesicle, fb=fin bud, pas=pharyngeal arches. Scale bar=100 microns.

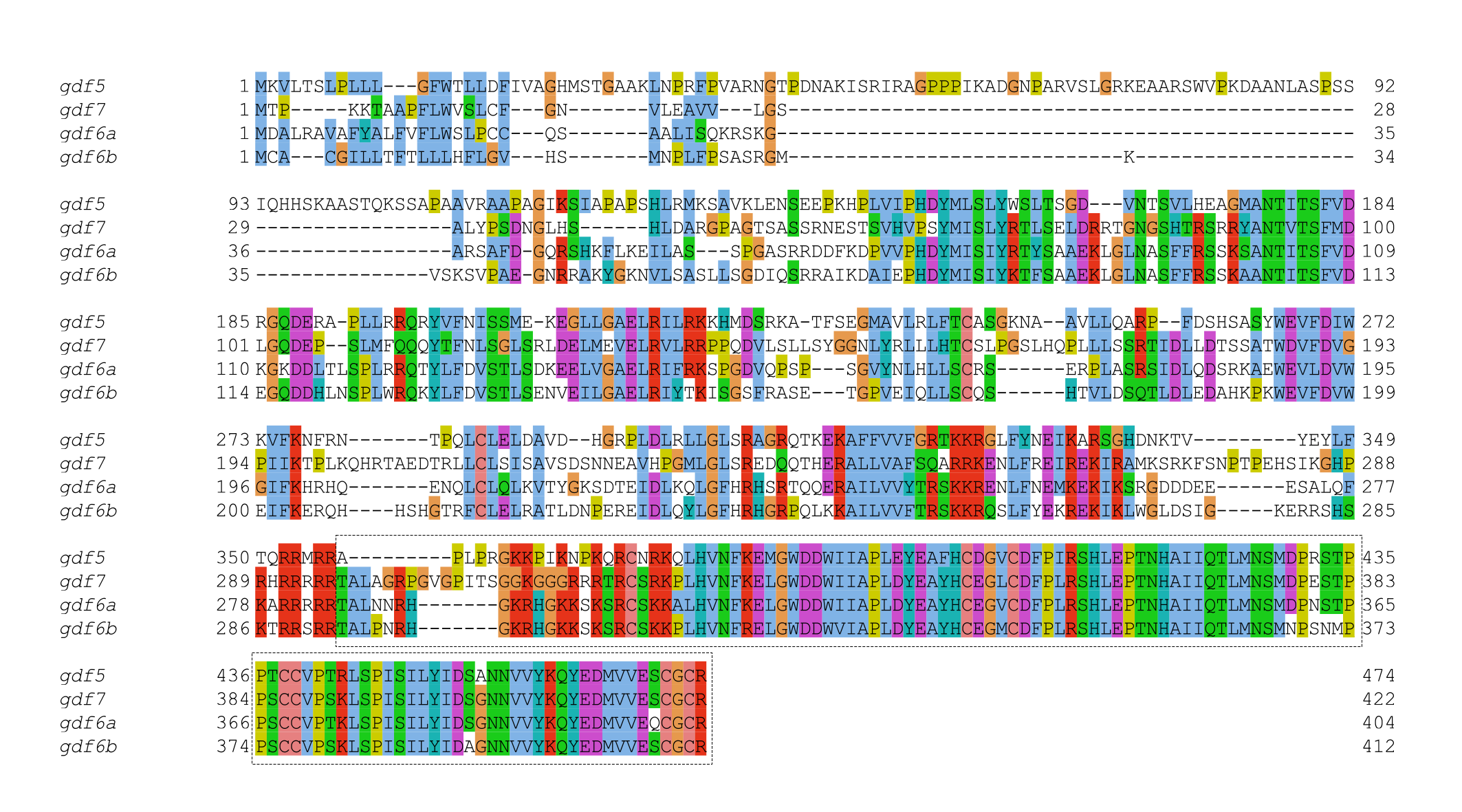

**Figure S2 - Multiple sequence alignment of Gdf5, Gdf6a, Gdf6b, and Gdf7 peptide sequences.** Colors correspond to amino acid residues that are conserved among all three polypeptide sequences, with each color corresponding to a specific amino acid. The sequence position is shown to the left and right of each amino acid sequence. Like their mammalian homologs, Gdf5, Gdf6, Gdf6b, and Gdf7 share a high degree of sequence similarity in the C-terminal region, which contains the mature, secreted signaling domain for each protein (indicated by the dotted box).

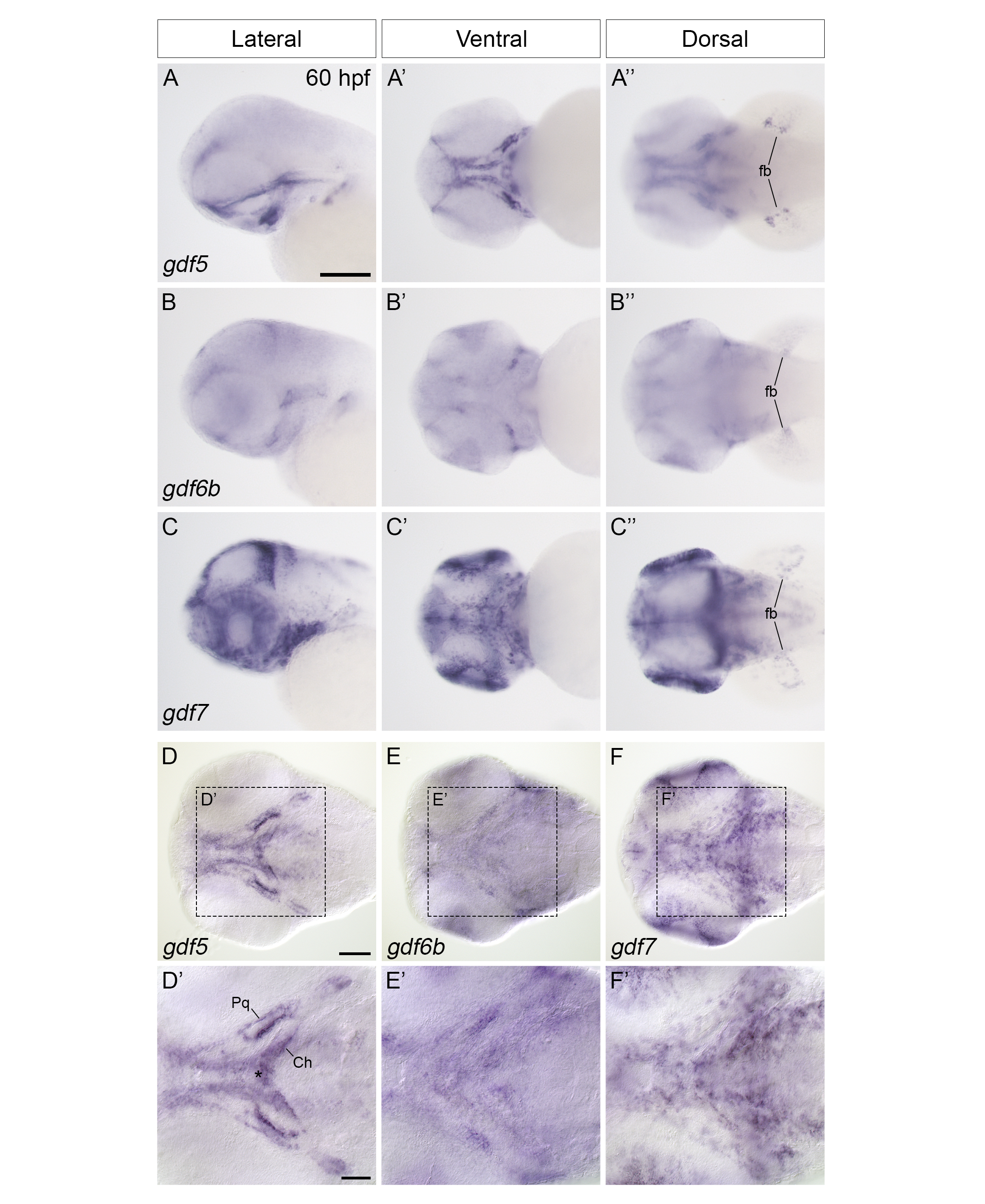

**Figure S3 – Expression of *gdf5*, *gdf6b*, and *gdf7*.** (A-C’’) *in situ* hybridization showing *gdf5* (A-A’’), *gdf6b* (B-B’’), and *gdf7* (C-C’’) mRNA localization in the head at 60 hpf. *gdf5* is localized to several skeletal tissues including the pharyngeal skeleton and neurocranium (A, A’) and the fin buds (A’’). *gdf6b* displays weak expression in the head (B, B’) and the fin buds (B’’). *gdf7* has widespread expression in the pharyngeal arches, brain, and eye (C-C’) and in the fin buds (C’’). (D-F’) DIC microscopy images of zebrafish embryos showing *gdf6a* mRNA expression in the ventral head at 60 hpf. *gdf5* expression is restricted to the perichondrum surrounding the palatoquadrate and ceratohyals (D, D’). *gdf5* is also expressed in the midline of pharyngeal arch 2 at the site of the presumptive basihyal and the joint that articulates the ceratohyals (D’, asterisk). *gdf6b* expression in the ventral head is diffuse and non-overlapping with *gdf6a* expression (E, E’). While *gdf7* expression in the ventral head is widespread, it is more superior than *gdf6a* expression and likely localized to the epidermis (F, F’). fb=fin bud, Pq=palatoquadrate, Ch=ceratohyal. Scale bars=100 microns in A and D and 50 microns in D’.

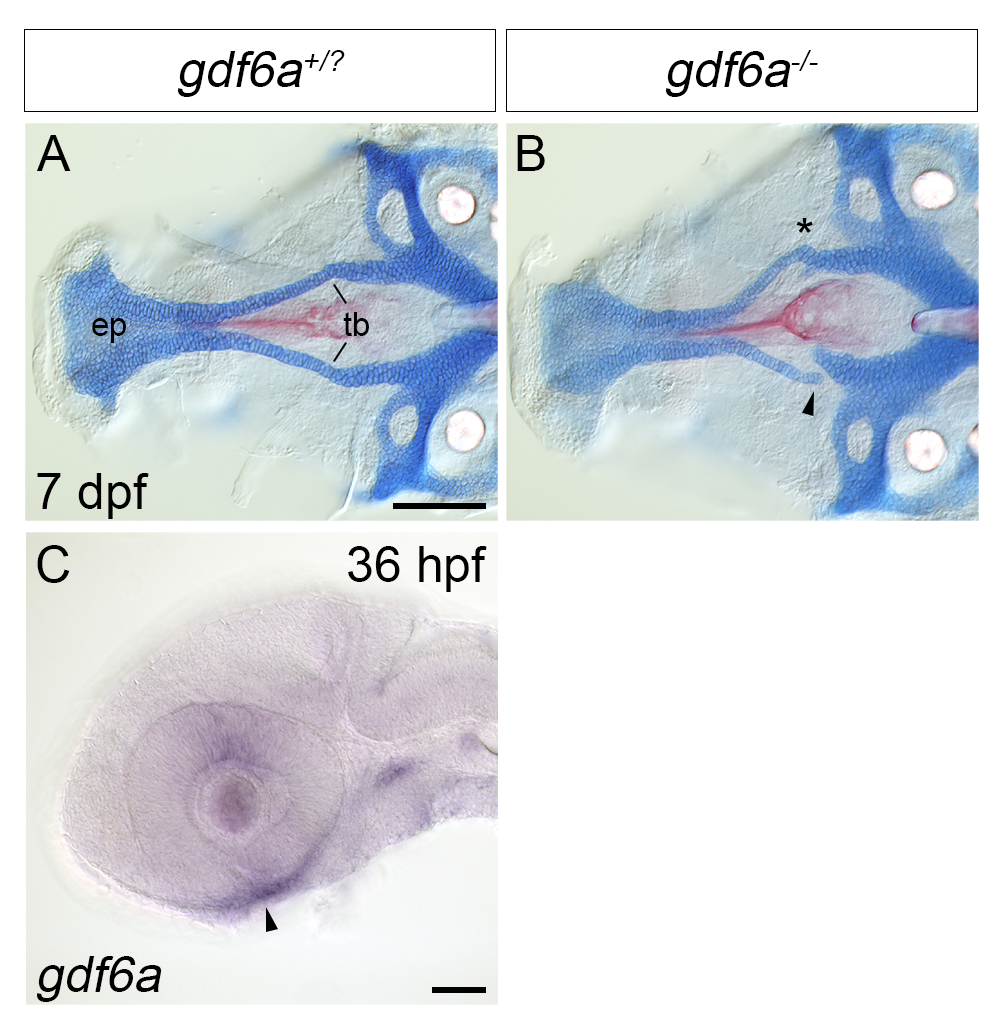

**Figure S4 – *gdf6a* regulates development of the neurocranium*.*** (A-B) DIC microscopy images of ventral flatmounts of neurocraniums from *gdf6a^+/?^* (A) and *gdf6a^-/-^* (B) larvae stained with Alcian blue and Alizarin red at 7 dpf. The overall length of the ethmoid plate is reduced in *gdf6a^-/-^* animals (B) compared to *gdf6a^+/?^* siblings (A). *gdf6a^-/-^* animals (n=19/19) also have defective trabeculae development; trabeculae frequently have gaps in them (B, arrowhead) or are bent and appear “fragile” (B, asterisk). 21% of larvae (4/19) have unilateral gaps in the trabeculae, 5% (1/19) have bilateral gaps, and 63% (12/19) have no gaps but trabeculae that are bent compared to *gdf6a^+/?^* siblings. (C) A DIC microscopy image of zebrafish embryos showing *gdf6a* mRNA expression in the head at 36 hpf. *gdf6a* is expressed in the mandibular domain at 36 hpf (arrowhead). ep=ethmoid plate, tb=trabeculae. Scale bars=100 microns in A, 50 microns in C.

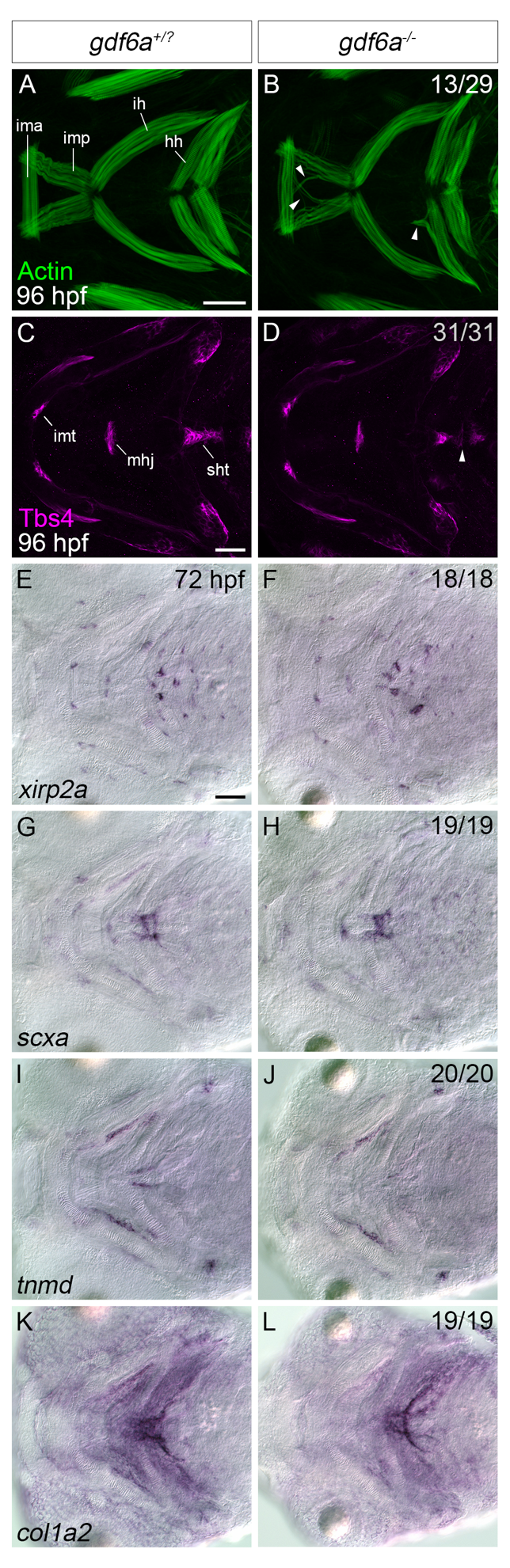

**Figure S5 – *gdf6a* regulates development of the craniofacial muscles independently of tendon formation*.*** (A-D) Single confocal z-slices of *gdf6a^+/?^* siblings (A, C) and *gdf6a^-/-^* mutant (B, D) at 96 hpf showing zebrafish craniofacial muscles (A, B) and tendons (C, D) visualized with phalloidin (for muscles) or Thrombospondin 4 (Tbs4) immunofluorescence (for tendons). *gdf6a^-/-^* mutants display the mislocalization of craniofacial muscle fibres at 96 hpf (B, n=13/29, arrowheads) compared to *gdf6a^+/?^* siblings (A). Specifically, the mislocalization takes place primarily in the intramandibularis anterior, the intramandibularis posterior, the interhyoideus, and hyohyoideus. This mislocalization occurs independently of tendon formation, as tendons form normally in *gdf6a^-/-^* mutants (D, n=31/31) compared to *gdf6a^+/?^* siblings (C). (E-L) DIC microscopy images showing the mRNA localization for the tendon markers *xirp2a* (E, F), *scxa* (G, H), *tnmd* (I, J), and *col1a2* (K, L) at 72 hpf in *gdf6a^+/?^* siblings *gdf6a^-/-^* mutants. All tendon markers show comparable expression patterns in *gdf6a^-/-^* mutants (n=18/18 for *xirp2a,* 19/19 for *scxa*, 20/20 for *tnmd,* and 19/19 for *col1a2*) compared to *gdf6a^+/?^* siblings. ima=intramandibularis anterior, imp=intramandibularis posterior, ih=interhyoideus, hh=hyohyoideus, imt=intermandibularis tendon, mhj=mandibulohyoid junction, sht=sternohyoideus tendon. Scale bar =50 microns.

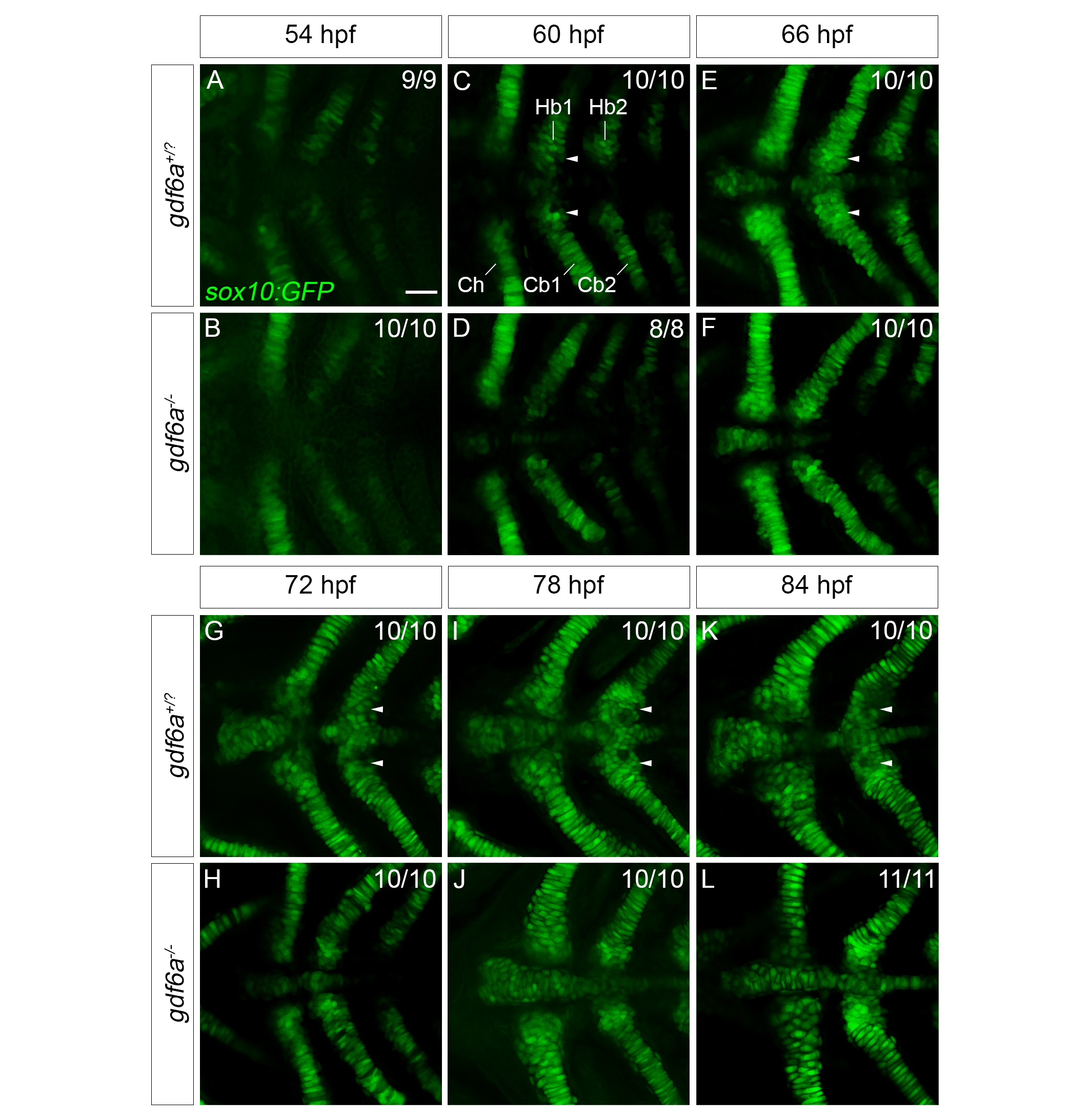

**Figure S6 – *gdf6a* regulates formation of the hypobranchials from 60 hpf onwards.** (A-L) Single confocal z-slices of the posterior pharyngeal arches of *gdf6a^+/?^;Tg(sox10:GFP)* and *gdf6a^-/-^;Tg(sox10:GFP)* zebrafish at 54, 60, 66, 72, 78, and 84 hpf. At 54 hpf, the pharyngeal cartilage has just condensed and started to differentiate in *gdf6a^+/?^* animals, as evidenced by the relatively weak *sox10:GFP* signal at this time and is comparable to *gdf6a^-/-^* mutant animals (n=10/10) (A-B). At 60 hpf, defects in hypobranchial formation become apparent in *gdf6a^-/-^* mutants (D) compared to *gdf6a^+/?^* siblings (C); while there are GFP+ cells corresponding to the hypobranchials present in *gdf6a^+/?^* siblings, these cells are absent in *gdf6a^-/-^* mutants (n=8/8). The absence of the hypobranchials in *gdf6a^-/-^* mutants becomes more apparent at 66 hpf; in *gdf6a^+/?^* siblings, GFP+ cells are beginning to aggregate to form the presumptive hypobranchials (E), which is defective in *gdf6a^-/-^* mutants (F, n=10/10). As development proceeds, it becomes apparent that the hypobranchials do not form in *gdf6a^-/-^* mutants (H, J) compared to siblings (G, I) at 72 and 78 hpf (n=10/10 for each age). By 84 hpf, *gdf6a^-/-^* mutants (n=11/11) have almost no GFP+ cells corresponding to the hypobranchials (L), which is in stark contrast to *gdf6a^+/?^* siblings (K), which have well-developed hypobranchials. Ch=ceratohyal, Hb1=hypobranchial 1, Hb2=hypobranchial 2, Cb1=ceratobranchial 1, Cb2=ceratobranchial 2. Scale bar=50 microns.

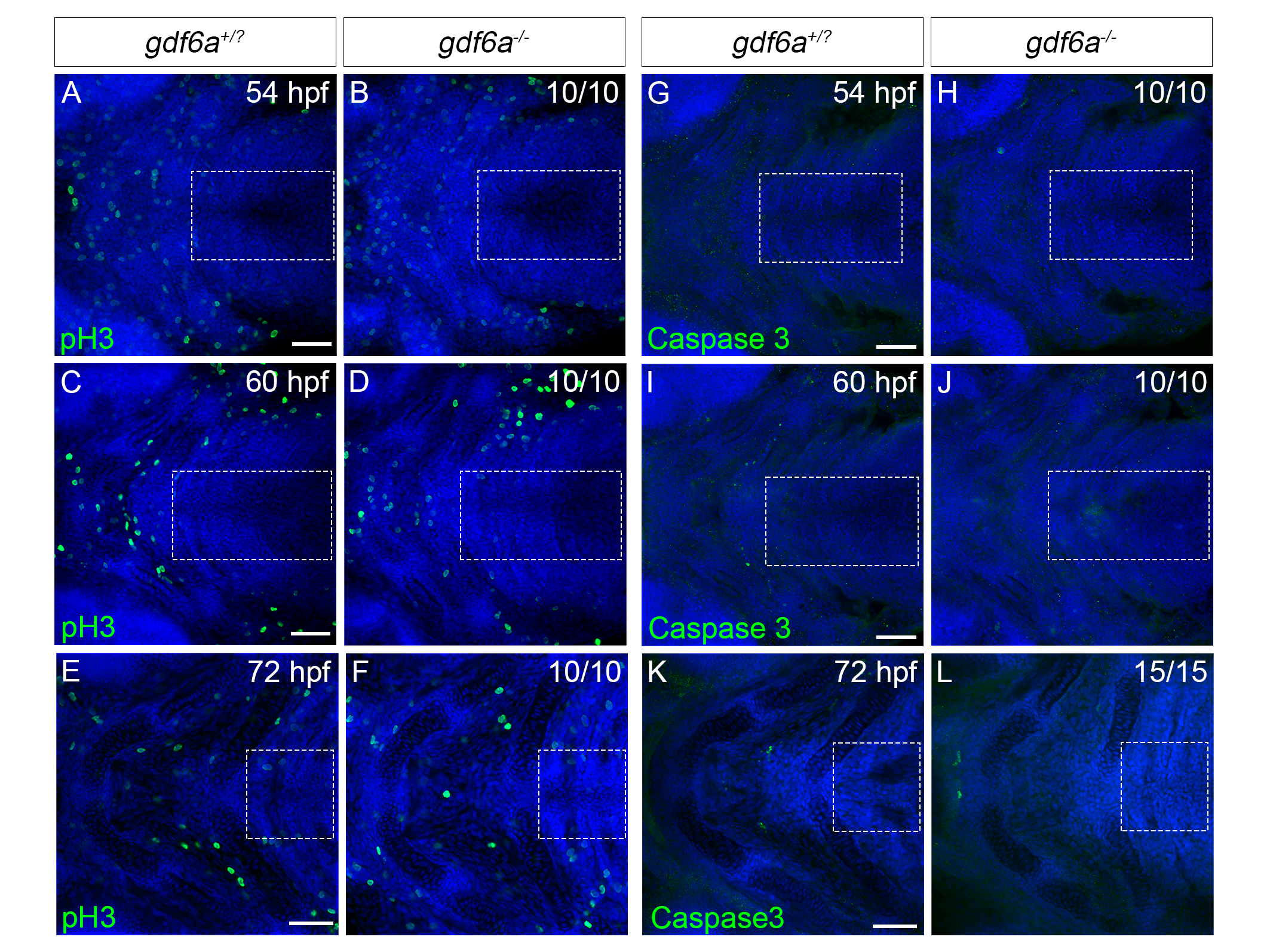

**Figure S7 – *gdf6a* does not regulate cell division or cell death in the posterior arches.** (A-F) Single confocal z-slices of *gdf6a^+/?^* and *gdf6a^-/-^* embryos at 54, 60, and 72 hpf showing the localization of phosphorylated Histone H3 (pH3) using immunofluorescence. At all stages, there are no pH3+ cells present in the posterior pharyngeal arches at any stage analyzed (A, C, E, dashed boxes). Similarly, *gdf6a^-/-^* mutants display no appreciable increase or decrease in pH3+ cells in the posterior pharyngeal arches (B, D, F, dashed boxes, n=10 for all stages). (G-L) Single confocal z-slices of *gdf6a^+/?^* and *gdf6a^-/-^* embryos at 54, 60, and 72 hpf showing the localization of cleaved Caspase 3 (Caspase 3) using immunofluorescence. At all stages, there are no Caspase 3+ cells present in the posterior pharyngeal arches at any stage analyzed (G, I, K, dashed boxes). Similarly, *gdf6a^-/-^* mutants display no appreciable increase or decrease in Caspase 3+ cells in the posterior pharyngeal arches (H, J, L, dashed boxes, n=10 for 54 and 60 hpf, n=15 for 72 hpf). Scale bars=50 microns.

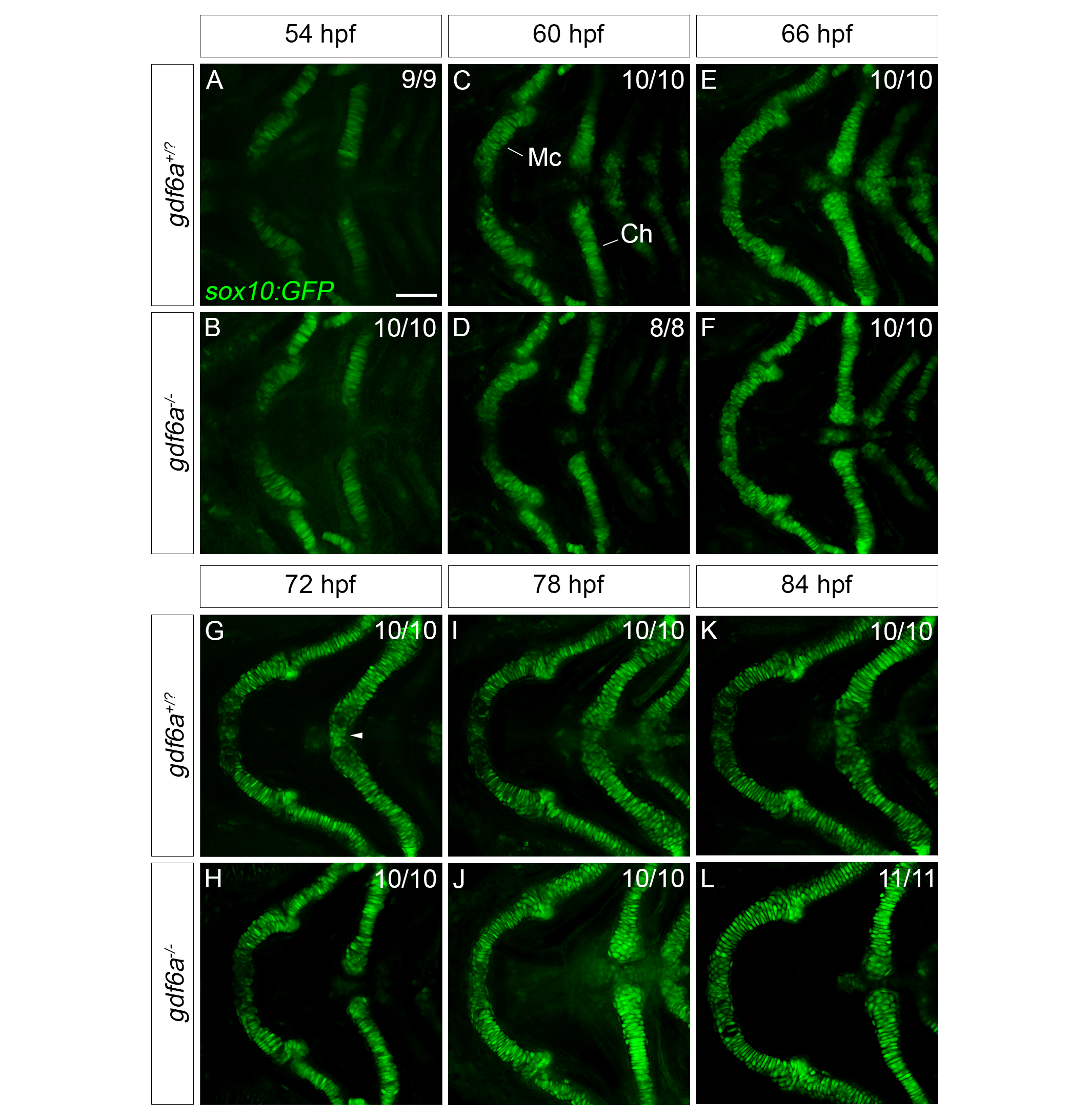

**Figure S8 – *gdf6a* regulates chondrogenesis of the ceratohyal joint from 72 hpf onwards.** (A-L) Single confocal z-slices of the mandibular and hyoid arches of *gdf6a^+/?^;Tg(sox10:GFP)* and *gdf6a^-/-^;Tg(sox10:GFP)* zebrafish at 54, 60, 66, 72, 78, and 84 hpf. From 54-66 hpf, the development of Meckel’s cartilage and ceratohyal are nearly indistinguishable between *gdf6a^-/-^* mutants (B, D, F, n=10/10, 8/8, 10/10 respectively and *gdf6a^+/?^* siblings (A, C, E). In contrast, from 72 hpf onwards, noticeable differences between *gdf6a^-/-^* mutants and *gdf6a^+/?^* siblings arise. In *gdf6a^+/?^* siblings, there is a bridge of chondrocytes at the ceratohyal joint that connects the bilateral arms of the ceratohyal (G, arrowhead); this chondrocyte bridge is missing in *gdf6a^-/-^* animals at the same stage (H, n=10/10). This results in the ceratohyals becoming progressively more misaligned at 78 hpf (J, n=10/10) and 84 hpf (L, n=11/11) in *gdf6a^-/-^* mutants compared to *gdf6a^+/?^* siblings at the same stages, where the ceratohyals are properly aligned (I, K).

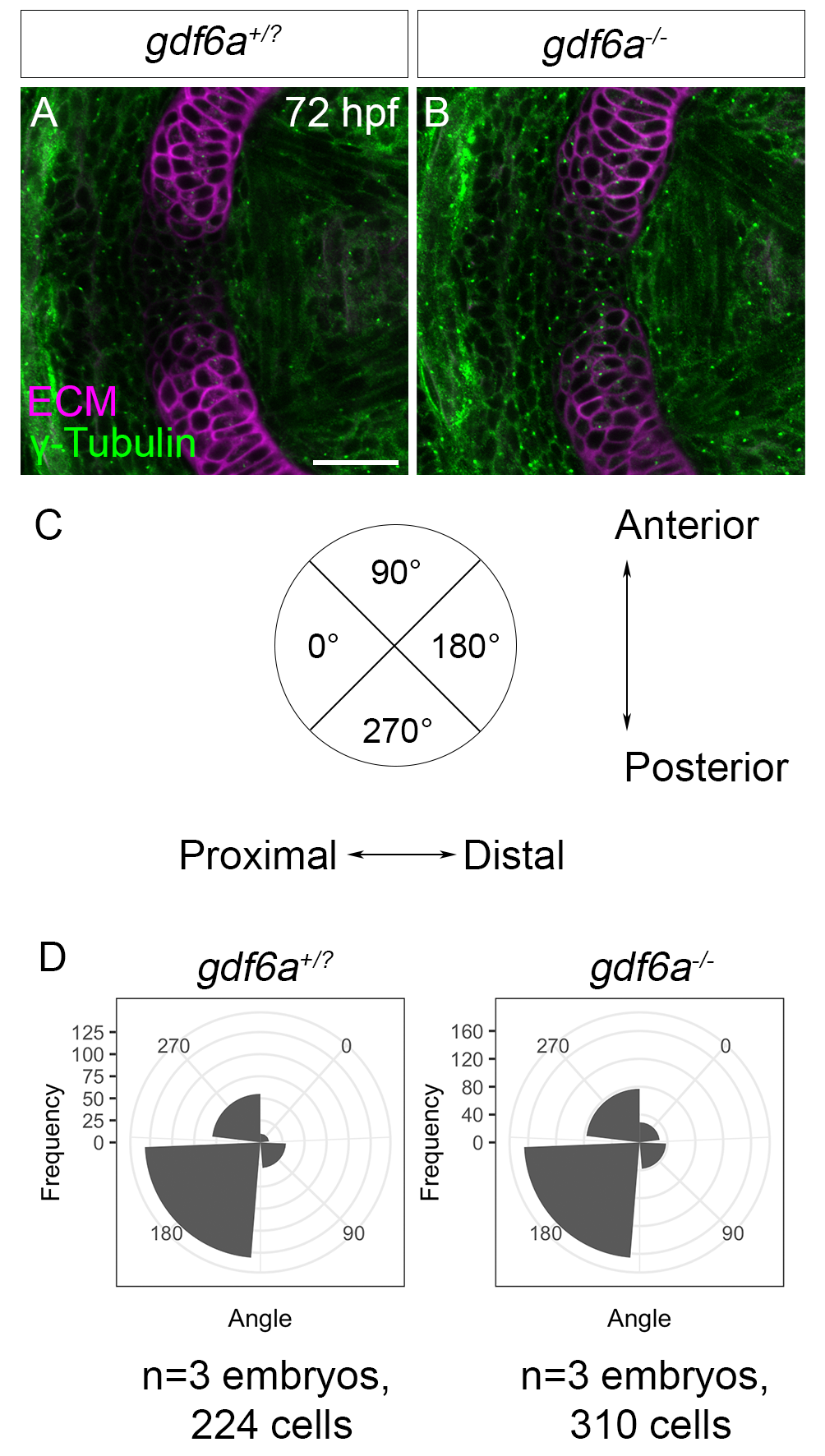

**Figure S9 – *gdf6a* does not regulate cell polarity in the mandibular symphysis.** (A-D) Single confocal z-slices of the mandibular symphysis from *gdf6a^+/?^* and *gdf6a^-/-^* animals at 72 hpf with the microtubule organizing center visualized with γ-Tubulin immunofluorescence and extracellular matrix visualized with wheat germ agglutinin. In *gdf6a^+/?^* animals, the microtubule organizing center tends to be oriented either toward or away from the midline of the mandibular symphysis (A), and this pattern is generally maintained in *gdf6a^-/-^* mutants (B). (C) Schematic showing how γ-Tubulin puncta were quantified. γ-Tubulin were sorted into bins based on their localization in the cell: puncta localized to the proximal side of the cell were designated as 0°, puncta at the anterior side of the cell were designated at 90°, puncta localized to the distal side of the cell were designated at 180° and puncta at the posterior side of the cell were designated as 270°. (D) Quantification of A and B. In *gdf6a^+/?^* animals, the MTOC was oriented on the proximal side in 9/224 cells, the anterior side of 29/224 cells, the distal side of 131/224 cells, and the posterior side of 55/224 cells (n=3 embryos, 224 cells). In *gdf6a^-/-^* animals, the MTOC was oriented on the proximal side in 29/310 cells, the anterior side of 38/310 cells, the distal side of 166/310 cells, and the posterior side of 77/310 cells (n=3 embryos, 224 cells). There is not a statistically significant difference in the distribution of the MTOC between *gdf6a^+/?^* and *gdf6a^-/-^* animals (Chi square test, χ^2^_3_=7.25, p=0.064). ECM=extracellular matrix. Scale bars=25 microns.

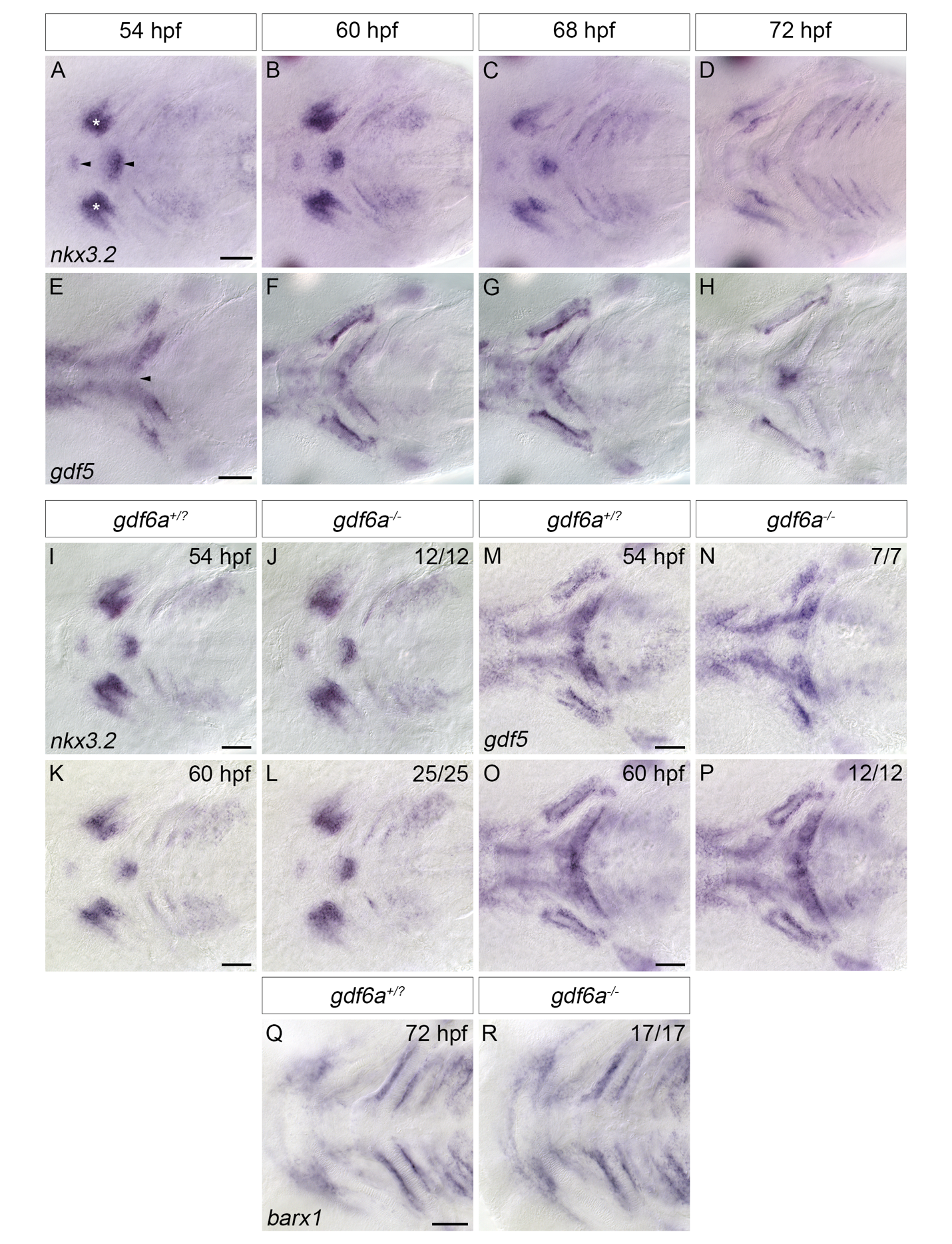

**Figure S10 – Joint patterning markers are unaffected in *gdf6a^-/-^* mutants.** (A-H) DIC microscopy images of zebrafish embryos showing *nkx3.2* and *gdf5* mRNA expression in the pharyngeal arches of wild type embryos from 54-72 hpf. At 54 hpf, *nkx3.2* is expressed at the site of the presumptive jaw joint (A, asterisks). There is also substantial *nkx3.2* expression in the presumptive mandibular symphysis and ceratohyal joint at this stage (A, arrowheads). This expression pattern persists until 60 hpf (B). By 68 hpf, the expression of *nkx3.2* in the mandibular symphysis and ceratohyal joint starts to fade (C), and by 72 hpf, there is little appreciable *nkx3.2 expression* in the mandibular symphysis and ceratohyal joint (D). At 54 hpf, *gdf5* mRNA is localized to the perichondrium of the ceratohyal and palatoquadrate (E, asterisks). At this stage, *gdf5* mRNA is also detected in the ceratohyal joint (E, arrowheads). The expression of *gdf5* in the perichondrium of the palatoquadrate and ceratohyal strengthens, as does its expression in the ceratohyal joint (F and G). At 72 hpf, the perichondral expression of *gdf5* has waned significantly, but expression in the ceratohyal joint is maintained (H). (I-L) DIC microscopy images of *gdf6a^+/?^* (I, K) and *gdf6a^-/-^* (J, L) zebrafish embryos showing *nkx3.2* mRNA expression at 54 (I, J) and 60 (K, L) hpf. The expression domain of *nkx3.2* in *gdf6a^-/-^* mutants (n=12/12 and n=25/25 for 45 and 60 hpf, respectively) (J, L) is almost indistinguishable from *gdf6a^+/?^* siblings (I, K) at both stages. (M-P) DIC microscopy images of *gdf6a^+/?^* (M, O) and *gdf6a^-/-^* (N, P) zebrafish embryos showing *gdf5* mRNA expression at 54 (M, N) and 60 (O, P) hpf. At 54 hpf, there is a gap separating the domain of *gdf5* expression in the ceratohyal joint in *gdf6a^-/-^* mutants (n=7/7) (N) compared to *gdf6a^+/?^* siblings (M). By 60 hpf, the expression of *gdf5* in *gdf6a^+/?^* siblings (O) is indistinguishable from *gdf6a^-/-^* mutants (P) (n=12/12). (Q-R) DIC microscopy images of *gdf6a^+/?^* (Q) and *gdf6a^-/-^* (R) zebrafish embryos at 72 hpf showing *barx1* mRNA expression at 72 hpf. In *gdf6a^+/?^* siblings, *barx1* is localized to the perichondrium of of Meckel’s cartilage and the ceratohyal and is largely absent from the developing mandibular symphysis and ceratohyal joint (A). In *gdf6a^-/-^* mutants (n=17/17), the expression of *barx1* mRNA is unchanged when compared to *gdf6a^+/?^* siblings, indicating that *gdf6a* does not regulate *barx1* expression during craniofacial development Scale bars=50 microns.

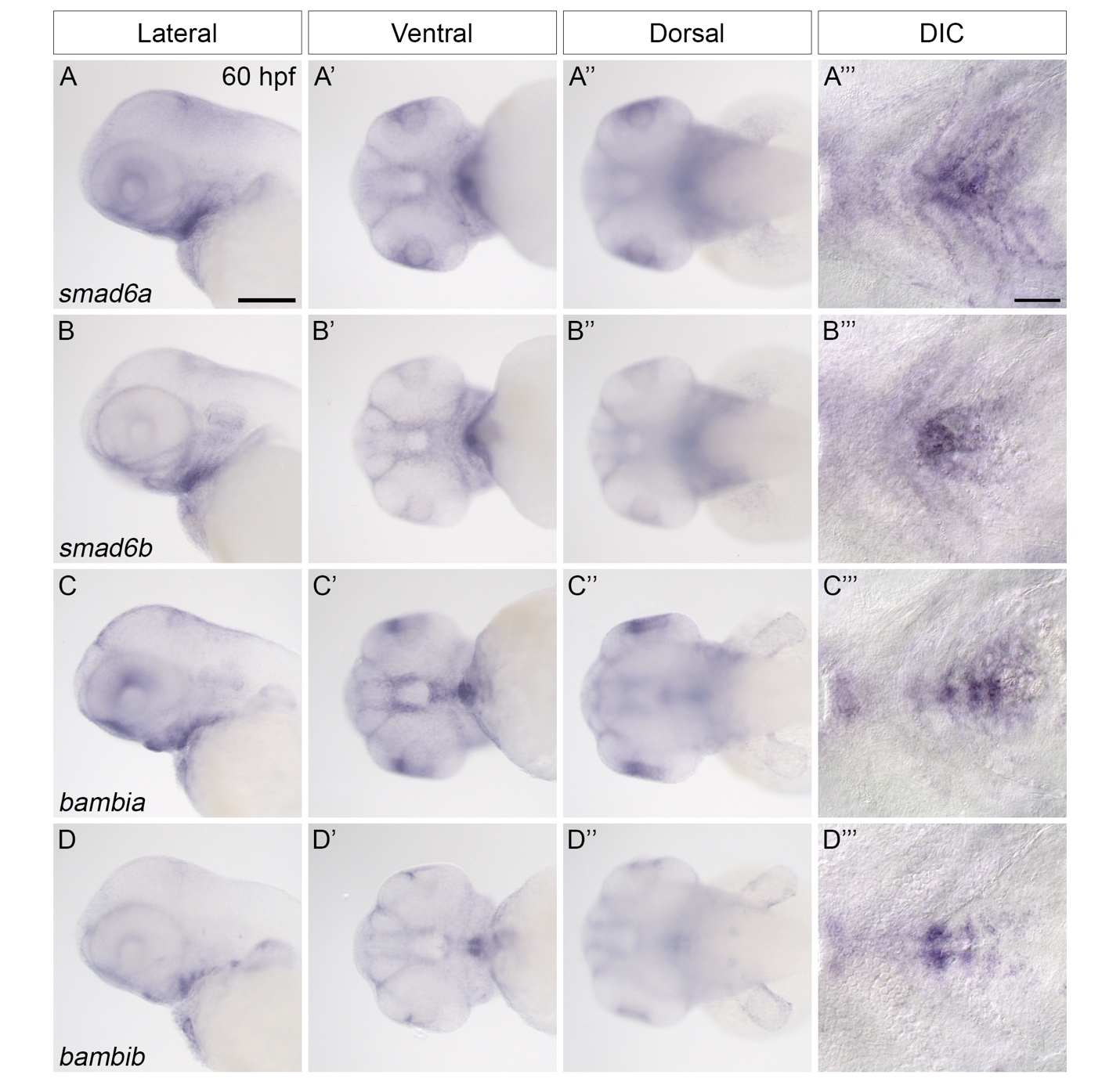

**Figure S11 – Expression of *smad6a, smad6b, bambia,* and *bambib*.** Whole mount dissecting microscope and DIC microscopy images of *in situ* hybridization showing *smad6a* (A-A’’’), *smad6b* (B-B’’’), and *bambia* (C-C’’’), *bambib* (D-D’’’), *nog1* (E-E’’’), mRNA localization in the head at 60 hpf. *smad6a, smad6b, bambia,* and *bambib* are all localized to the midline of the pharyngeal arches (A-D’’’). Scale bars=100 microns in A, 50 microns in A’’’.

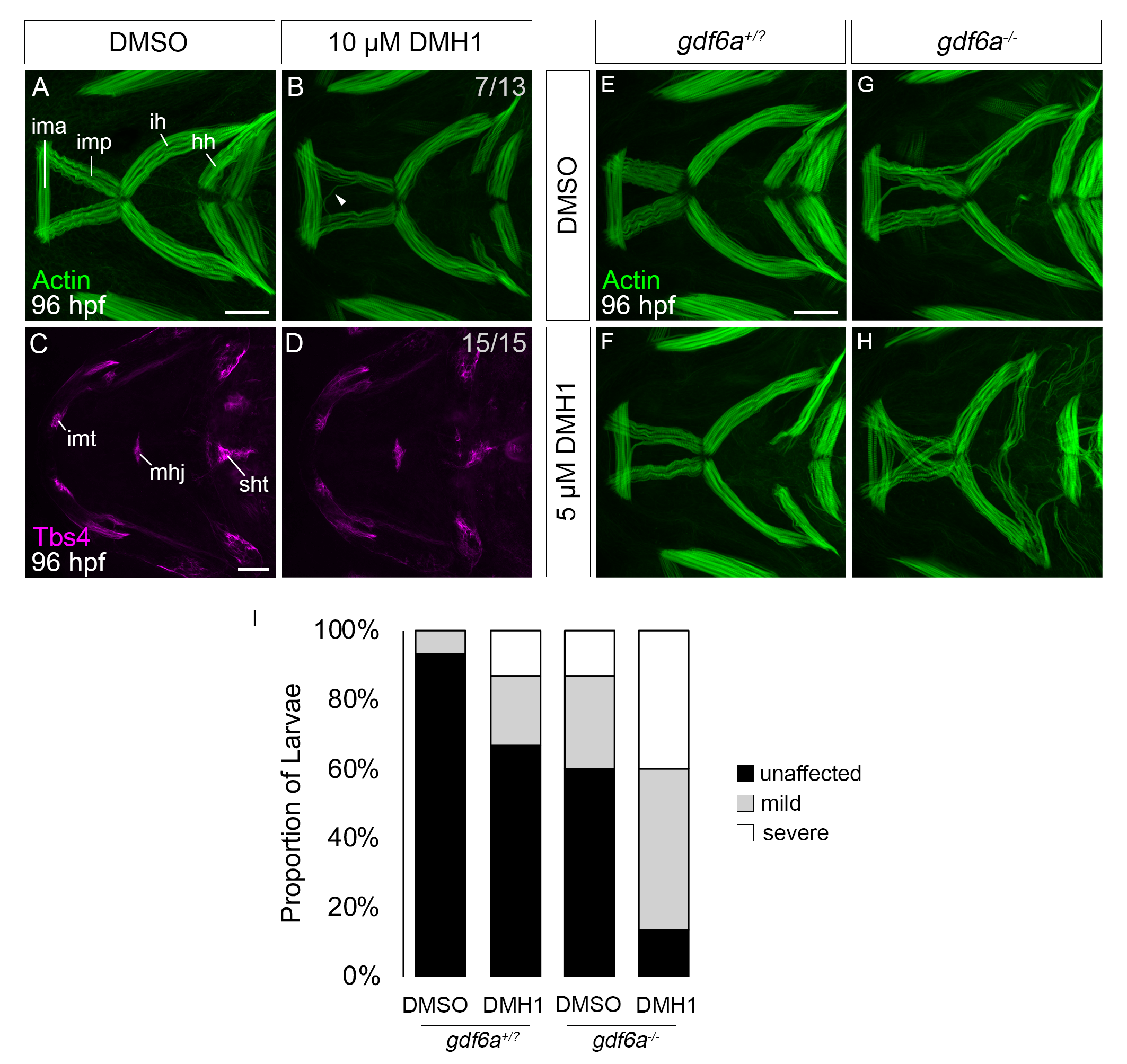

**Figure S12 – BMP signaling is necessary for proper craniofacial muscle development.** (A-D) Maximum intensity projections of craniofacial muscles (A, B) or tendons (C, D) from zebrafish treated with DMSO (A, C) or 10 μM DMH1 (B, D) from 48-72 hpf. Muscles were visualized with phalloidin (for muscles) and tendons were visualized with Thrombospondin 4 (Tbs4) immunofluorescence (for tendons) at 96 hpf. Similar to what is observed in *gdf6a^-/-^* mutants, several DMH1-treated larvae display (n=7/13) the mislocalization of craniofacial muscle fibers (B, arrowhead) compared to DMSO-treated controls (A). This mislocalization occurs independently of tendon formation, as craniofacial tendons develop normally in all DMH1-treated animals (n=15/15, D) compared to DMSO-treated controls (C). Results are representative of N=2 independent experiments. (E-H) Maximum intensity projections of craniofacial muscles from *gdf6a^+/?^* (E, F) and *gdf6a^-/-^* (G, H) larvae treated with DMSO (E, G) or 5μM DMH1 (F, H) from 48-72 hpf. When treated with DMSO, almost all *gdf6a^+/?^* siblings (n=14/15) show normal craniofacial muscle localization at 96 hpf. In contrast, *gdf6a^-/-^* mutants (n=5/15) larvae show either mild (n=3/15) or severe (n=2/15) mislocalization of muscle fibers (G). When treated with 5 μM DMH1, *gdf6a^+/?^* siblings display either a mild (n=4/15) or severe (2/15) muscle fiber mislocalization (F), similar to what is observed when *gdf6a^-/-^* mutants are treated with DMSO (G). Almost all (n=13/15) *gdf6a^-/-^* animals treated with DMH1 display either mild (n=7/15) or severe (n=6/15) muscle fiber localization (H). (I) Quantification of the results in E-H. Craniofacial muscles from individual larvae were sorted into the following groups and their proportion was quantified: Unaffected (similar to what is displayed in E), mildly affected (similar to what is shown in F and G), and severely affected (similar to what is shown in H). Results are representative of N=2 independent experiments. ima=intramandibularis anterior, imp=intramandibularis posterior, ih=interhyoideus, hh=hyohyoideus, imt=intermandibularis tendon, mhj=mandibulohyoid junction, sht=sternohyoideus tendon. Scale bars=50 microns.

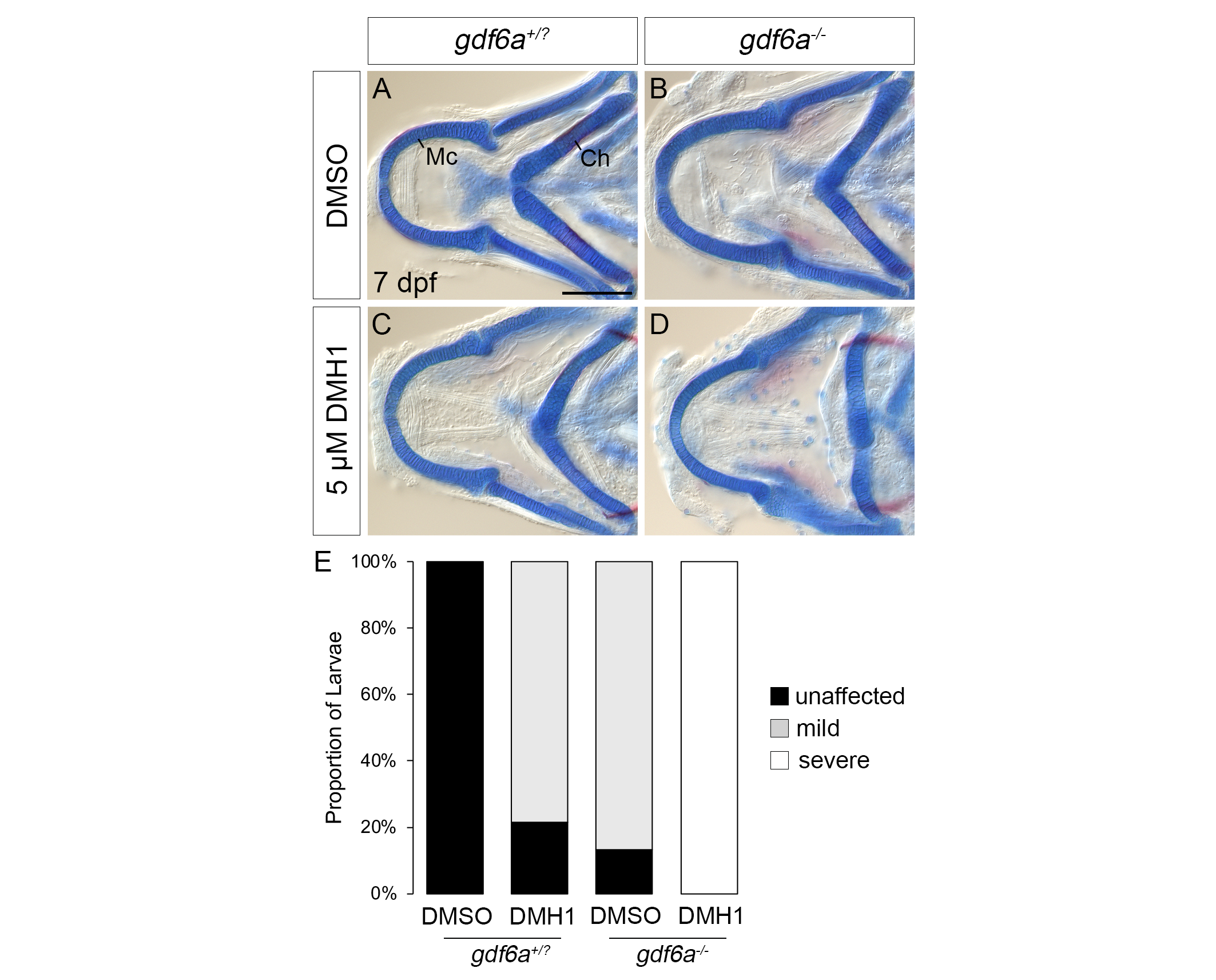

**Figure S13 – mutations in *gdf6a* synergize with a low dose of DMH1.** (A-D) DIC microscopy images of flatmounted 7 dpf larval craniofacial cartilage visualized with Alcian blue and Alizarin red from *gdf6a^+/?^* and *gdf6a^-/-^* larvae treated with DMSO or 5μM DMH1 from 48-72 hpf. Almost all *gdf6a^+/?^* (A, n=11/12) and *gdf6a^-/-^* (B, n=14/15) larvae treated with DMSO show no phenotype and a relatively mild phenotype, respectively. Most *gdf6a^+/?^* (C, n=11/14) treated with DMH1 display a mild phenotype similar to DMSO-treated *gdf6a^-/-^* animals. In contrast, all *gdf6a^-/-^* animals treated with DMH1 (D, n=11/11) display a phenotype that is more severe than any other condition. (E) Quantification of the results in E-H. The jaws from individual larvae were sorted into the following groups and their proportion was quantified: Unaffected (similar to what is displayed in A), mildly affected (similar to what is shown in B and C), and severely affected (similar to what is shown in D). Results are representative of N=2 independent experiments. Mc=Meckel’s cartilage, Ch=ceratohyal. Scale bar=100 microns.

**Table S1** – Sequences of primers used to make riboprobes for wholemount *in-situ* hybridization. The T7 promoter site is underlined.

| Primer name | Sequence (5'-3') |
| --- | --- |
| *gdf6a* probe F | CAGTCGCCTTTTACGCGCTC |
| *gdf6a* probe R T7 | TAATACGACTCACTATAGGGGGCGATGATCCAGTCGTCCCA |
| *gdf5* probe F | CACCCCCGATAACGCGAAGA |
| *gdf5* probe R T7 | TAATACGACTCACTATAGGGGGTACTCCAGCGGAGCGATGA |
| *gdf6b* probe F | GTTTCCAAGCGCATCCAGGG |
| *gdf6b* probe R T7 | TAATACGACTCACTATAGGGGCTGAGTTTGGAGGGGACGCA |
| *gdf7* probe F | CTCCCAAGAAGACTGCCGCT |
| *gdf7* probe R T7 | TAATACGACTCACTATAGGGGCAACTGGGAGGGGTGGACTC |
| *nog3* probe F | TATTTCCTGGCCACCGTGCT |
| *nog3* probe R T7 | TAATACGACTCACTATAGGGGACAAATGAATCAGGCACCACACT |
| *smad6a* probe F | GAAAACTGGTGCCGTGACCG |
| *smad6a* probe R T7 | TAATACGACTCACTATAGGGGCCTCACACTGTTCGGGTCGT |
| *smad6b* probe F | GGACCAAGATGGGGGTACGG |
| *smad6b* probe R T7 | TAATACGACTCACTATAGGGGCCGTCTAGCAGCTCCGACTC |
| *bambia* probe F | TCGCCTGGTTTCTCTGTGGTT |
| *bambia* probe R T7 | TAATACGACTCACTATAGGGGTGCAGTCCATAAGCTCCACCC |
| *bambib* probe F | CAGCGGAACAGCAGTCTCGT |
| *bambib* probe R T7 | TAATACGACTCACTATAGGGGCCAGCATAAAGCCGCTCCTC |
| *sox9a* probe F | GTTTATGGTGTGGGCGCAGG |
| *sox9a* probe R T7 | TAATACGACTCACTATAGGGAGGGCACCCCTGTAGAGTCA |
| *col2a1a* probe F | TGATTCGGGGACTGTGCTGT |
| *col2a1a* probe R T7 | TAATACGACTCACTATAGGGTACCAGGGTTACCGGATGCAC |
| *nkx3.2* probe F | CGAGCTGGATGTGTGTTTCT |
| *nkx3.2* probe R T7 | TAATACGACTCACTATAGGGCTCCTCAAGTCCAGCAAATGT |
| *barx1* probe F | TTTGGAGATTGGGGCGCACT |
| *barx1* probe R T7 | TAATACGACTCACTATAGGGAATCACCACTCGGTCCCTGG |

**Table S1** – Sequences of oligonucleotides used to generate DNA templates for gRNA synthesis.

| Oligonucleotide name | Sequence (5'-3') |
| --- | --- |
| smad5_Target_1 | TAATACGACTCACTATAGGTTACCATTCCCAGGTCGCGTTTTAGAGCTAGAAATAGC |
| smad5_Target_2 | TAATACGACTCACTATAGGTCGTGGCACCAGAACCGGGTTTTAGAGCTAGAAATAGC |
| smad5_Target_3 | TAATACGACTCACTATAGGTATCAGGAGCCCAGTCACGTTTTAGAGCTAGAAATAGC |
| smad5_Target_4 | TAATACGACTCACTATAGGTAGTTGCAGTTTCGACTCGTTTTAGAGCTAGAAATAGC |
